# Effect of *phyB* and *phyC* loss-of-function mutations on the wheat transcriptome under short and long day photoperiods

**DOI:** 10.1101/2020.04.07.030197

**Authors:** Nestor Kippes, Carl VanGessel, James Hamilton, Ani Akpinar, Hikmet Budak, Jorge Dubcovsky, Stephen Pearce

## Abstract

**Background:** Photoperiod signals provide important cues by which plants regulate their growth and development in response to predictable seasonal changes. Phytochromes, a family of red and far-red light receptors, play critical roles in regulating flowering time in response to changing photoperiods. A previous study showed that loss-of-function mutations in either *PHYB* or *PHYC* result in large delays in heading time and in the differential regulation of a large number of genes in wheat plants grown in an inductive long day (LD) photoperiod.

**Results:** We found that under non-inductive short-day (SD) photoperiods, *phyB*-null and *phyC*-null mutants were taller, had a reduced number of tillers, longer and wider leaves, and headed later than wild-type plants. Unexpectedly, both mutants flowered earlier in SD than LD, the inverse response to that of wild-type plants. We observed a larger number of differentially expressed genes between mutants and wild-type under SD than under LD, and in both cases, the number was larger for *phyB* than for *phyC*. We identified subsets of differentially expressed and alternatively spliced genes that were specifically regulated by *PHYB* and *PHYC* in either SD or LD photoperiods, and a smaller set of genes that were regulated in both photoperiods. We observed significantly higher transcript levels of the flowering promoting genes *VRN-A1*, *PPD-B1* and *GIGANTEA* in the *phy-*null mutants in SD than in LD, which suggests that they could contribute to the earlier flowering of the *phy-*null mutants in SD than in LD.

**Conclusions:** Our study revealed an unexpected reversion of the wheat LD plants into SD plants in the *phyB*-null and *phyC*-null mutants and identified candidate genes potentially involved in this phenomenon. Our RNA-seq data provides insight into light signaling pathways in inductive and non-inductive photoperiods and a set of candidate genes to dissect the underlying developmental regulatory networks in wheat.

## Background

As sessile organisms, plants must be able to respond to fluctuations in their environment to maximize their reproductive success. To achieve this, plants have evolved a series of regulatory mechanisms to ensure that critical stages of their development coincide with optimal environmental conditions. One important determinant of reproductive success is flowering time, which is strongly influenced by seasonal changes in photoperiod and temperature [1]. In cereal crops, these cues are fundamental to ensure the plant does not flower too early, to prevent exposure of sensitive reproductive tissues to late-spring frosts, or too late, so as to minimize exposure to damaging high temperatures during grain filling [2]. There is a direct link between reproductive success and grain production, so characterizing the regulatory networks underlying flowering time is critical to support the development of resilient crop varieties, to help meet the world’s growing demand for food [2].

Plants respond differently to seasonal variation in photoperiod according to the environment to which they are adapted. Whereas some plant species exhibit accelerated flowering in short day photoperiods (SD plants), others flower more rapidly in long days (LD plants). A third class of plants are day-neutral and flower irrespective of the photoperiod. The temperate cereals, including common wheat (*Triticum aestivum* L.), are LD plants. This ensures that plants remain in a vegetative phase during winter until the lengthening days of spring trigger the irreversible transition to reproductive development [1]. An additional requirement for a long period at low temperatures (vernalization) prevents flowering during the fall, when the days are still relatively long [3].

In wheat and other temperate cereals, the length of the night, rather than the length of the day, is critical for the perception of inductive photoperiods. This has been demonstrated by experiments in which exposing wheat plants to night-breaks (15 m periods of light in the middle of a long night) for at least 12 d was sufficient to accelerate flowering [4]. Loss-of-function mutations in the wheat phytochrome genes *PHYTOCHROME B* (*PHYB*) or *PHYC*, or in the *PHOTOPERIOD1* (*PPD1*) gene abolish the acceleration of flowering by night-breaks, suggesting that these genes are critical to measure the duration of the night [4].

A recent study in *Brachypodium* proposed a mechanism for the role of these genes in the determination of the photoperiodic response [5]. Phytochromes, a class of red (R, ~650 nm) and far-red (FR, ~720 nm) light receptors exist as one of two interchangeable forms, P_R_ and P_FR_. In darkness, the biologically inactive P_R_ form accumulates in the cytoplasm, but upon absorption of R light, P_R_ is converted to the bioactive P_FR_ form and is translocated to the nucleus [6-8]. Conversely, exposure to FR light causes the rapid reversion of P_FR_ to the P_R_ form, a reaction that also takes place more gradually during the night (dark or thermal reversion). Therefore, the duration of the night affects the amount of the bioactive P_FR_ form, which has been proposed to be critical for the degradation of the clock protein EARLY FLOWERING 3 (ELF3), a direct repressor of *PPD1* [5]. High ELF3 protein levels and the repression of *PPD1* have been proposed as the main cause of the late flowering phenotypes of the *phyC* mutant in *Brachypodium* [5].

*PPD1* encodes a PSUEDO-RESPONSE REGULATOR (PRR)-family protein that acts as a positive regulator of flowering in the LD grasses [9-11] but as a LD-repressor in the SD grasses rice [12] and sorghum [13], where this gene is referred to as *PRR37*. In wheat, allelic variation at the *PPD1* locus affects photoperiod sensitivity. Whereas the wild-type *Ppd-A1b* allele is expressed at very low levels during the night, the *Ppd-A1a* allele, which carries a promoter deletion encompassing the ELF3 binding site, shows increased transcript levels during the day and, particularly, at night [14]. Wheat varieties that carry the *Ppd-A1b* allele are referred to as photoperiod sensitive (PS) and those that carry *Ppd-A1a* as photoperiod insensitive (PI) because they exhibit accelerated heading under SD and reduced differences in heading time between SD and LD. It is important to point out that wheat varieties carrying the PI allele still show a significant acceleration of heading under LD [9, 11]. *PPD1* induces the expression of *FLOWERING LOCUS T1* (*FT1*), which encodes a protein with similarity to the PEBP family [15]. The FT1 protein is translocated through the phloem to the shoot apical meristem, where it forms a hexameric floral activation complex that directly activates the expression of meristem identity genes including *VERNALIZATION 1* (*VRN1*) and *FRUITFULL 2* (*FUL2*). These MADS-box genes play critical roles in triggering reproductive development [16-18]. In the cereals, *ft1*-null mutants exhibit a strong delay in flowering [19].

In addition to their role in the regulation of ELF3, bioactive P_FR_ phytochromes interact in the nucleus directly with PHYTOCHROME INTERACTING FAMILY (PIF) proteins, a class of bHLH transcription factors [20, 21]. In Arabidopsis, these interactions induce biochemical changes in the PIF proteins, which result in their ubiquitination and degradation via the 26S proteasome pathway [22]. In this species, accumulating PIF proteins act primarily as negative regulators of light signaling transcriptional networks, so their degradation in response to R light triggers a cascade of photoperiod-mediated transcriptional responses. Despite its important role in the light signaling pathway in Arabidopsis, the role of PIF proteins in the regulation of the photoperiod response in the temperate cereals remains unknown.

Phytochromes can also induce transcriptional variation through modulating alternative splicing (AS) [23]. In Arabidopsis, changes in AS were detected in over 1,500 genes in response to R light, in a PHYB-dependent manner [23]. These target genes include PIF3, whereby greater levels of P_FR_ PHYB increased the frequency of an intron retention event in this gene, disrupting the translated protein’s function [24]. In the moss *Physcomitrella patens*, the phytochrome protein PpPHY4 interacts directly with a splicing regulator to mediate AS in response to light [25]. Previously, the splicing factor RRC was found to mediate phytochrome response in Arabidopsis, suggesting this mechanism may be conserved in angiosperms [26].

Monocot genomes contain three phytochrome genes, *PHYA, PHYB* and *PHYC*, with three homeologous copies of each gene in hexaploid wheat [27]. In wheat and *Brachypodium*, both *PHYB* and *PHYC* are required for timely flowering in LD conditions and plants carrying non-functional copies of either phytochrome exhibit extreme delays in flowering, as well as changes in their vegetative morphology [28-30]. Using *phyB*-null and *phyC*-null Ethyl-methane sulfonate (EMS)-derived mutants in the tetraploid wheat variety ‘Kronos’, we previously described the sets of genes regulated by *PHYB* and *PHYC* in LDs [31]. Despite similar delays in flowering time in both mutants, we found that *PHYB* regulates approximately six times as many genes as *PHYC*, and that only a small core of 104 genes were regulated by both phytochromes at the transcriptional level [31]. These commonly regulated genes include several well-characterized flowering time genes, such as *PPD1* and *FT1*, and meristem identity genes, including *VRN1* and *FUL2*.

The role of the wheat phytochromes in non-inductive photoperiods remains an open question. Previously, we found that while *phyC*-null mutants flower later than WT plants in both SD and LD photoperiods, the effect is approximately five-fold smaller in SDs [28]. There is a significant interaction between photoperiod and *PHYC*, with the wild-type plants heading earlier in LDs than in SDs, and the *phyC*-null mutants heading earlier in SDs than LDs [28]. In the current study, we found that *phyB*-null Kronos mutants also flower significantly earlier in SD than in LD.

To characterize the genes involved in the earlier heading of the *phyB*-null and *phyC*-null mutants in SDs than in LDs, we compared the transcriptomes of these mutants under SD and LD conditions. We identified sets of genes regulated by *PHYB* and *PHYC* in both SD and LD photoperiods, as well as genes that were regulated only under a specific photoperiod. In addition, we found that both *PHYB* and *PHYC* regulate alternative splicing events in a number of genes, of which only a small proportion also showed significant differences in transcript levels between wild-type and *phy* mutants. The findings of this study contribute to our understanding of the complex regulatory networks controlling photoperiod-mediated flowering in wheat.

## Results

### Effect of *phyB*-null and *phyC*-null mutants on heading time

We first characterized the effect of Kronos-*phyB-*null and Kronos-*phyC-*null mutants on heading time under LD and SD conditions relative to wild-type Kronos (WT), a photoperiod insensitive (*Ppd-A1a*) spring wheat (*Vrn-A1*). The wild-type Kronos headed at 47 d in LD and at 95 d in SD (48 d delay, *P* < 0.0001), as expected for a LD plant. This result showed that Kronos plants carrying the *Ppd-A1a* allele still respond to changes in photoperiod. By contrast, both *phyB*-null and *phyC*-null mutants headed earlier in SD than in LD (108 d earlier for *phyB*-null, *P* < 0.001, Figure 1a, 23 d earlier for *phyC*-null, *P* < 0.001, Figure 1b). This reversal was the result of a much larger delay in heading time in the null mutants under LD (104 d and 196 d later than WT) than under SD (31 d and 39 d later than WT, *P* <0.0001, Figure 1a-b). The interactions between photoperiod and genotype were significant for both *PHYB* and *PHYC* (Figure 1a-b, *P* < 0.0001) [28].

**Figure 1:**
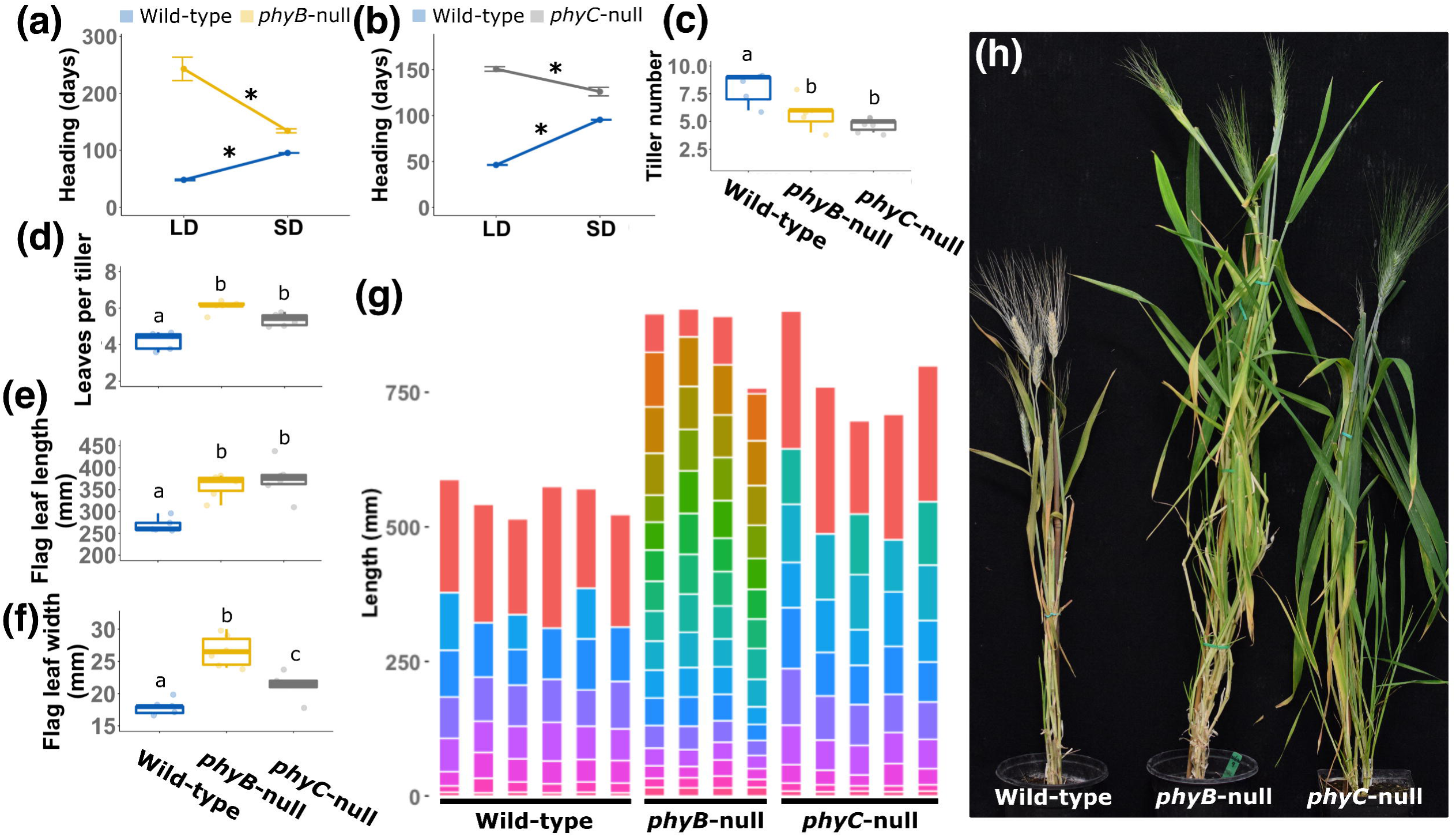
Phenotypic characterization of *phyB*-null and *phyC*-null mutants under SD conditions (8h light/16h dark). **(a)** Heading date of wild-type and *phyB*-null plants in SD and LD showing the significant interaction between *PHYB* and photoperiod**. (b)** Heading date of wild-type and *phyC*-null plants in SD and LD showing the significant interaction between *PHYC* and photoperiod. **(c)** Tiller number per plant. **(d)** Mean leaf number per tiller. **(e)** Flag leaf length. **(f)** Flag leaf width. **(g)** Internode length and number. Each bar represents an individual plant and the horizontal lines correspond to the position of the nodes. Each internode is represented by a different color, ordered according to their position in the stem. The uppermost segment in each individual represents the length between the last node and the spike (peduncle) **(h)** Picture of representative plants when *phyB*-null plants reached heading date. (**c** to **f**) Boxplots represent values of at least five biological replications. Different letters indicate significant differences (Tukey’s test *P* < 0.05). For (a) and (b), * signifies significant differences between photoperiods for each genotype, *P* < 0.0001. The differences between wild-type and mutant alleles were also highly significant (*P* < 0.0001) for both genes and both photoperiods.

Kronos plants carrying a single null allele in either the A or B homeologs of *PHYB* or *PHYC* showed no significant delay in heading date relative to the WT (Additional file 1, Figure S1) and the same was observed for other traits, so all subsequent results describe comparisons between *phyB*-null, *phyC*-null mutants and the WT in a Kronos-PI background.

### Effect of phyB-null and phyC-null mutants on plant phenotype under SD

We next characterized the vegetative phenotypes of these mutant lines under SD conditions. Tiller number was significantly lower in both mutants compared to the WT (Figure 1c), while mean leaf number per tiller was significantly higher in both mutants than in WT plants (Figure 1d), likely due to the delayed transition of the shoot apical meristem to the reproductive phase. In both *phyB*-null and *phyC*-null mutants, flag leaves were significantly longer and wider than WT (Figure 1e-f).

Stem development was also affected in the *phy* mutants. Both mutants were significantly taller than WT plants (*phyB*-null 310 mm taller, *P* = 9.72^E-06^ and *phyC*-null 220 mm taller, *P* = 0.00016, Figure 1g). While the *phyB*-null and *phyC*-null mutants did not differ significantly from one another in overall height, their stem structure was markedly different. The *phyB*-null mutants exhibited a larger number of internodes than either WT (9 more internodes than WT, *P* = 7.14^E-09^) or *phyC*-null mutants (7 more internodes than *phyC*, *P* = 3.53 ^E-07^), while *phyC*-null plants had a slightly increased internode number compared to the WT control (2.1 more internodes, *P* = 0.00013, Figure 1g). Representative plants of each genotype are shown in figure 1h, which was taken when *phyB*-null mutants reached heading date. Taken together, these results show that both *PHYB* and *PHYC* play important roles in regulating vegetative and reproductive development in non-inductive SD conditions.

### Characterizing the *PHYB-* and *PHYC-*regulated wheat transcriptome under SD

To investigate the transcriptional changes associated with the earlier flowering of the *phyB*-null and *phyC*-null plants relative to WT in the Kronos background, we performed an RNA-seq experiment in WT, *phyB*-null and *phyC*-null plants under SD conditions. We collected tissue from the last fully expanded leaf of four biological replicates per genotype at eight-weeks of age (Additional file 1, Figure S2). To facilitate comparison with a previous RNA-seq study of the same materials in LD conditions [31], we took samples at the same point of the photoperiod (four hours after dawn). We harvested tissues from eight-week-old plants in our SD experiment so the wild-type plants were at a similar developmental stage as the wild-type plants in the LD RNA-seq study, which were sampled at four-weeks of age.

After trimming raw reads for quality and adapter contamination, an average of 45.0 M trimmed 100 bp single-end reads per sample were mapped to unique positions in the IWGSC RefSeq v1.0 genome assembly (Additional file 1, Table S1). Using all normalized read counts mapped to high and low confidence gene models for each sample, we generated a multi-dimensional scaling (MDS) plot (Figure 2a). Samples grouped into three distinct clusters according to their genotype, reflecting consistent differences in overall transcriptome profile between genotypes and limited differences among biological replicates (Figure 2a).

**Figure 2:**
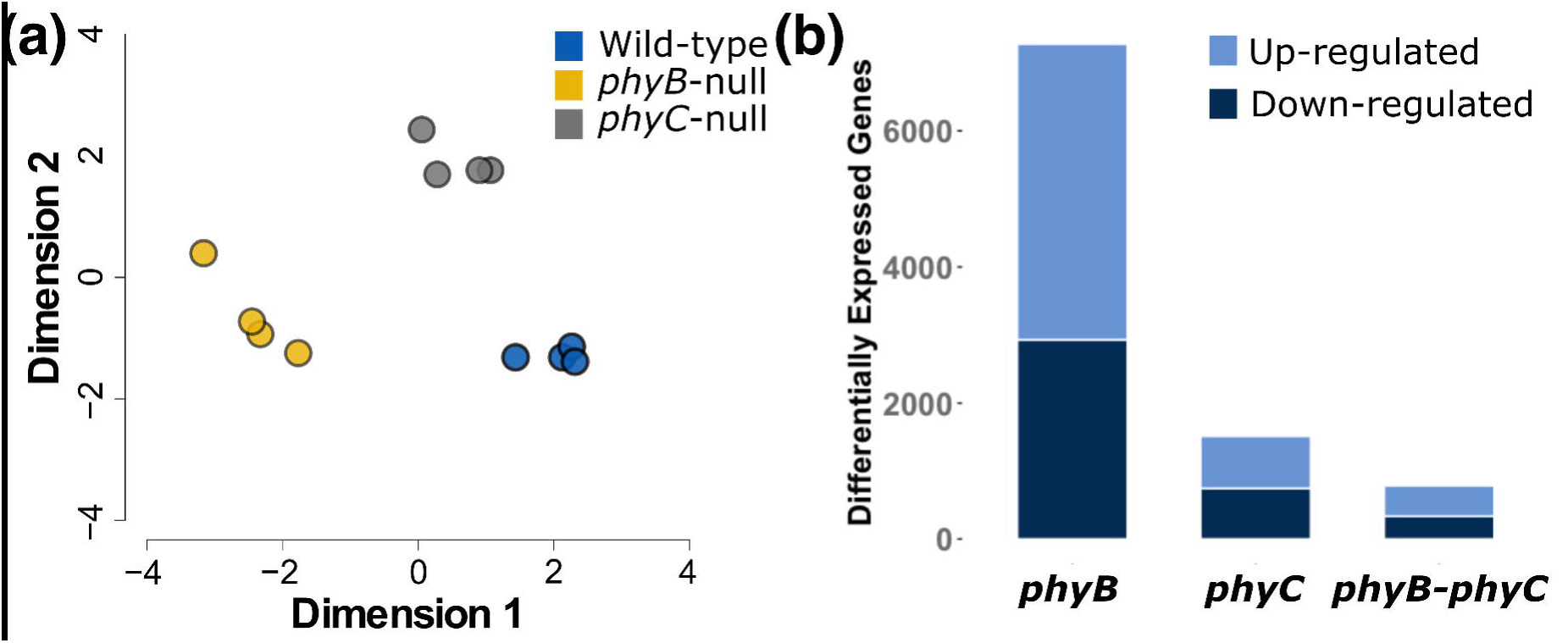
Transcriptomes of WT, *phyB*-null and *phyC*-null plants under SD photoperiods. **(a)** Multi-dimensional scaling (MDS) plot showing overall transcriptome profile of four biological replicates of each genotype. **(b)** Number of differentially expressed genes from pairwise comparisons between WT and *phyB*-null, WT and *phyC*-null and the subset of genes commonly regulated by both genes. Note that 27 additional genes were regulated by both *PHYB* and *PHYC* but in opposite directions and are not included in this graph.

We next performed pairwise comparisons between WT and both mutants to identify *PHYB-* and *PHYC*-differentially expressed (DE) genes under SD conditions. We found that 4.8 times as many genes were regulated by *PHYB* (7,272 DE genes) than by *PHYC* (1,511 DE genes, Figure 2b). Among these DE genes, a greater proportion were positively regulated by *PHYB* (59.7% with higher expression in WT than *phyB*-null) than by *PHYC* (50.6% of genes). There were 815 genes regulated by both *PHYB* and *PHYC*, including 783 genes regulated in the same direction and 27 in the opposite direction (upregulated by *PHYB* and downregulated by *PHYC* or vice versa, Figure 2b). Full details of expression data and statistical tests for each pairwise comparison are provided in Additional file 2.

To identify putative functions associated with these transcriptional changes, we performed a GO enrichment analysis for each subset of differentially expressed genes. Among the 7,272 genes regulated by *PHYB* in SDs, the most significantly enriched terms included ‘oxidation-reduction process’ and ‘protein phosphorylation’, while among the 1,511 genes regulated by *PHYC*, significant terms included ‘defense response’ and ‘cellular iron homeostasis’ (Additional file 1, Table S2). In genes commonly regulated by both *PHYB* and *PHYC*, enriched terms included ‘defense response’ and ‘protein phosphorylation’ (Additional file 1, Table S2).

Changes in development are often associated with differential expression of genes encoding transcription factors. Compared to the overall proportion of genes encoding transcription factors in our dataset (3.2% of 72,120 expressed genes), an increase was observed for the *PHYB*-(5.3%) and *PHYC*-regulated genes (5.4%), and an even larger increase was detected among the genes regulated by both *PHYB* and *PHYC* (6.5%). More importantly, several critical genes involved in the regulation of flowering were differentially expressed in *phyB* and *phyC* relative to the WT. Transcript levels of *PPD-B1*, both homeologs of *FT1* and *FT2*, *VRN-B1* and *SOC1* were all significantly lower in both *phyB*-null and *phyC*-null mutants compared to WT plants (Additional file 2).

To validate these expression data and to study longer-term trends of the expression of these genes in SD conditions, we performed qRT-PCR analysis for selected candidate genes across six time points, using the same genotypes as in the RNA-seq analysis. At the eight-week time point, the qRT-PCR experiment confirmed the RNA-seq results, showing that transcript levels of *VRN1*, *FT1*, *FT2*, *PPD1* and *FT3* were all significantly lower in *phyB*-null and *phyC*-null mutants compared to WT (Figure 3). *PPD1* expression was significantly higher in WT than either mutant at all assayed time points. It is important to note that these values represent the combined transcript levels of *Ppd-A1a* and *Ppd-B1b* homeologs.

**Figure 3:**
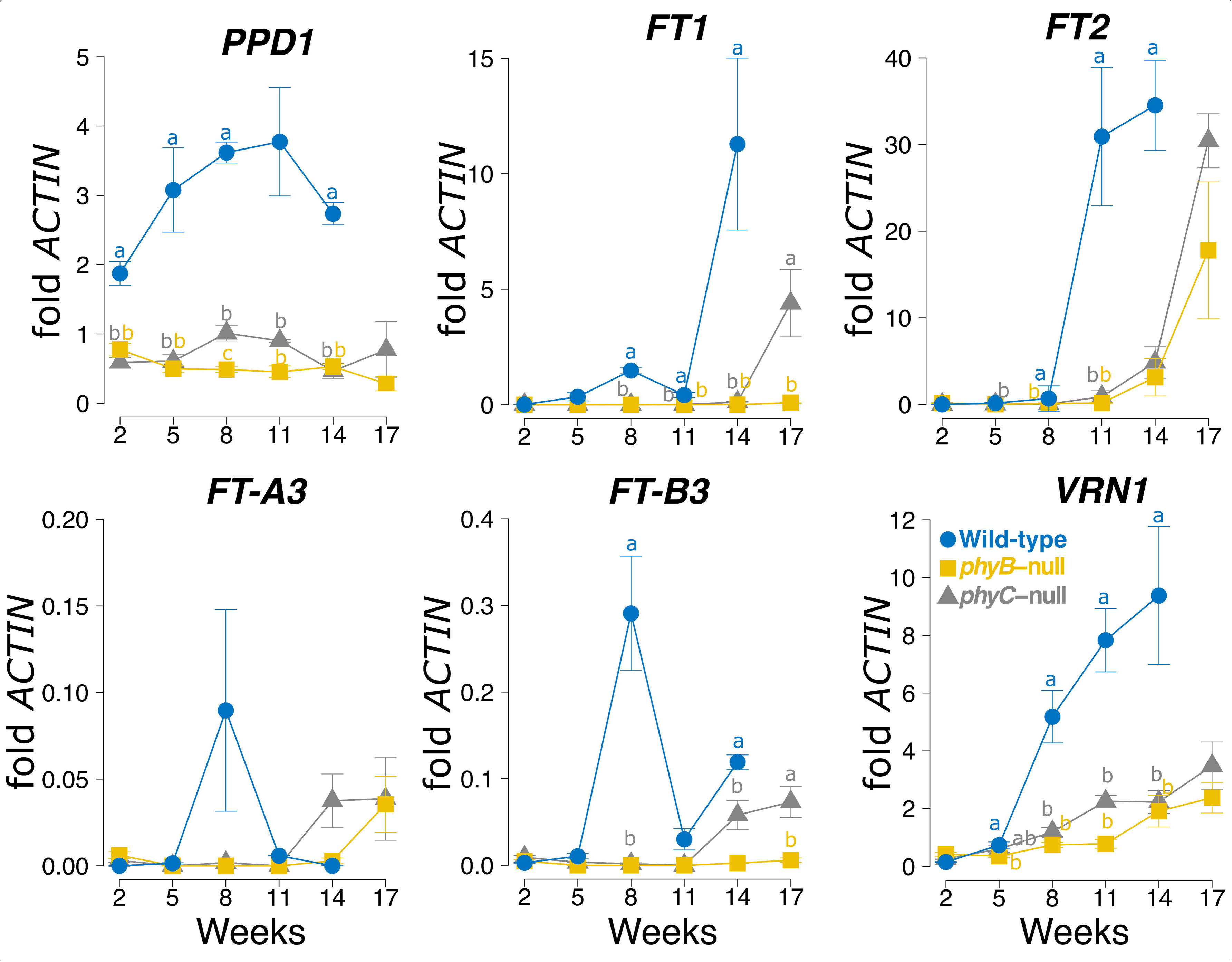
Transcript levels of flowering time genes in WT, *phyB*-null and *phyC*-null mutants under SD conditions assayed by qRT-PCR. Each data point represents the mean of four biological replications and error bars represent SEM. Different letters denote significant differences between samples at the 0.05 confidence level. All primers used to assay expression were redundant for A and B homeologs, except for *FT-A3* and *FT-B3*. The WT control headed at 14 w, and at 17 w plants showed signs of senescence so were not sampled.

There were also differences in the expression profiles of members of the *FT-like* family between genotypes. In wild-type plants, *FT1* transcript levels were more than double the levels of *FT2* at the 5 w and 8 w time points (Figure 3), consistent with results from a previous study [32]. By contrast, in both *phyB* and *phyC* plants, *FT2* was upregulated at an earlier time point (14 w) than *FT1*, which increased in expression at 17 w in *phyC* mutants, but remained low throughout the experiment in *phyB* mutants (Figure 3). *FT3* was expressed at much lower levels than *FT1* and *FT2*, and in the wild-type both *FT-A3* and *FT-B3* showed a transient peak in expression at 8 w. In the *phyC* mutant, *FT3* levels started to increase at 14 w and were even higher at 17 w, whereas in the *phyB* mutant we only observed upregulation of *FT-A3* at 17 w (Figure 3). *VRN1* expression increased gradually in both mutant lines throughout this time course, but its transcript levels remained significantly lower than in WT lines at all time points from 5 w onwards (Figure 3).

Taken together, these differences in expression between phytochrome mutants and WT are consistent with the delayed heading date of *phyB* and *phyC* mutants compared to WT. These results also confirm that phytochromes play an important role in the regulation of critical flowering genes under both SD and LD photoperiods. The earlier expression of *FT2* relative to *FT1* and its high transcript levels (Figure 3), suggest that this gene may play a more important role in the early flowering of the phytochrome mutants under SD than under LD.

### Effect of photoperiod on phytochrome-regulated genes

To explore the effect of photoperiod on the differences between WT and the phytochrome mutants, we compared the DE genes generated in the current study in SD collected from 8-week-old plants, with the DE genes in a previous dataset that used the same plant materials grown in LD conditions collected from 4-week-old plants [31]. This SD time point was chosen to match developmental stage in the WT plants between SD and LD conditions (Waddington stage 3 [33]).

To allow a direct comparison between datasets, we remapped the RNA-seq reads from our earlier LD study to the IWGSC RefSeq v1.0 genome assembly using the same mapping and quantification parameters adjusted for read length. Using this updated genomic reference, 52.8% of all reads mapped uniquely (Additional file 1, Table S3). This LD dataset includes two experimental replicates, each with four biological replications. Genes were considered differentially expressed only when significant in both experiments. This approach reduces the false positive rate, but means that direct comparisons of the number of differentially expressed genes between SD and LD datasets should be approached with caution because SD data represents only a single experimental replicate.

An MDS-plot separating the samples on the basis of their whole transcriptomic profiles revealed a high consistency between experimental replicates, but wider differences between genotypic classes (Additional file 1, Figure S3). We identified 3,668 genes that were differentially expressed between WT and *phyB*-null mutants in both experimental replicates and 424 genes for the corresponding comparisons with the *phyC*-null mutant. Just 141 of these genes were regulated by both *PHYB* and *PHYC* under LD conditions. With slight variations, these results are consistent with our previous study mapping these sequencing data to an older version of the wheat genome [31]. Full details of expression data and statistical tests for each pairwise comparison in LD photoperiods are provided in Additional File 3.

In Figure 4, we divided genes into mutually exclusive classes according to the conditions under which they were differentially expressed between wild-type and mutant alleles (i.e. regulated by *PHYB* or *PHYC* under either SD or LD conditions). For clarity, this figure excludes some pairwise comparisons with low numbers of genes, so the numbers presented in the text do not sum to the complete number of DE genes, which are presented in Additional File 4. In both photoperiods, a greater number of genes were regulated only by *PHYB* than only by *PHYC* (Figure 4). In SDs, 9.6-fold more genes were specifically regulated by *PHYB* (5,369 genes) than *PHYC* (561 genes), whereas in LDs, 13.5-fold more genes were specifically regulated by *PHYB* (2,289 genes) than by *PHYC* (167 genes, Figure 4).

**Figure 4:**
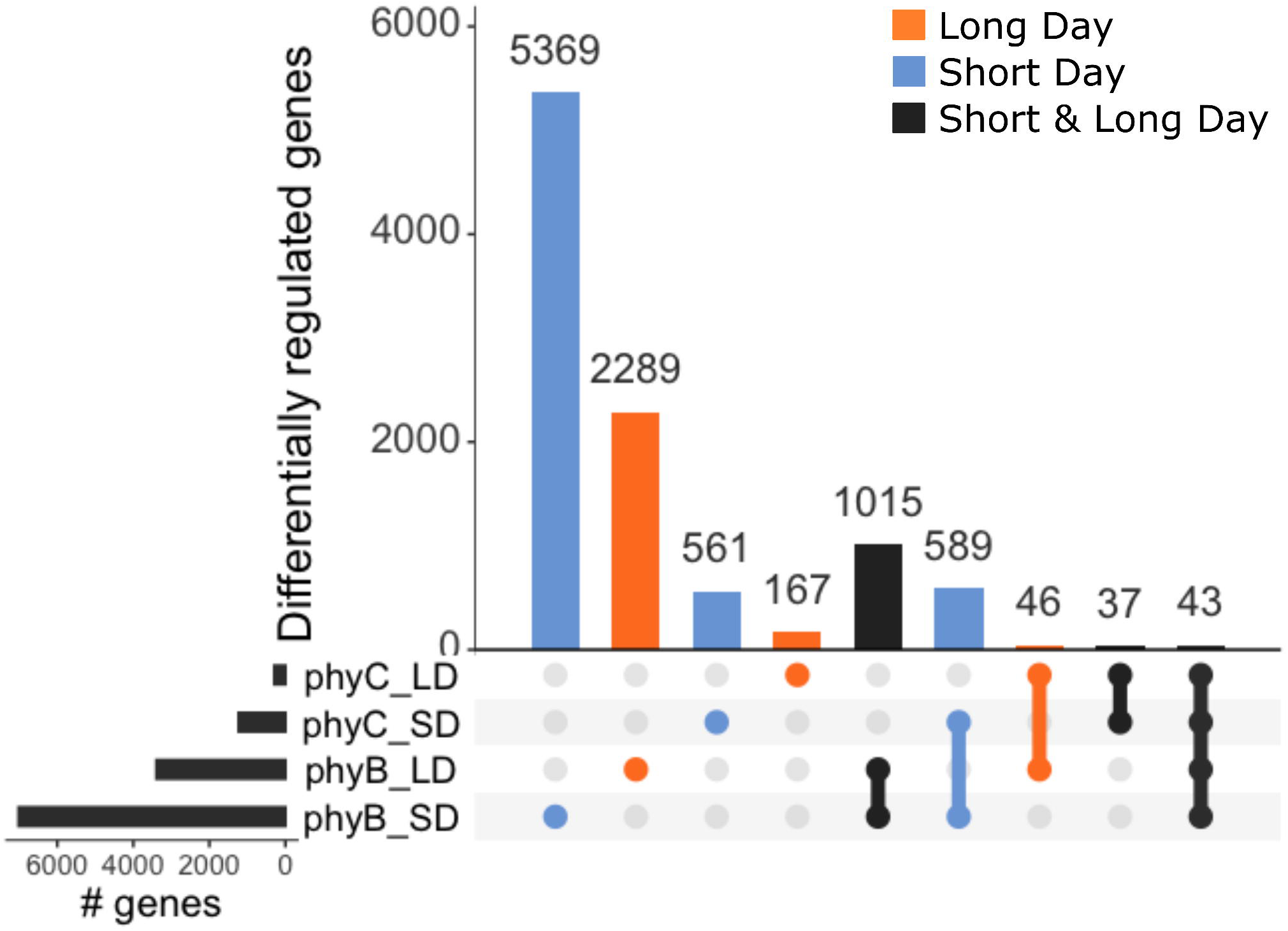
Summary of differentially expressed genes regulated by *PHYB* and *PHYC* in either SD or LD photoperiods. Each mutually exclusive category includes genes differentially expressed between WT and the respective phytochrome mutant in pairwise comparisons. For clarity, not all pairwise comparisons presented in Additional File 4 are displayed here.

There were more genes differentially expressed between WT and mutant genotypes exclusively in SD (589 genes) than exclusively in LD (46 genes, Figure 4). In addition, the number of genes differentially regulated in a single photoperiod was larger than the number of genes differentially regulated in both photoperiods. For example, there were 1,015 genes regulated by *PHYB* in both SD and LD, compared to 5,369 and 2,289 genes that were significant in either SD or LD photoperiods, respectively (Figure 4). Since the LD acceleration of heading time in wheat requires the presence of both *PHYB* and *PHYC*, we focused on genes DE in both mutants. We detected 589 of these DE genes in SD only, 46 in LD only and 43 in both SD and LD (Figure 4).

In the GO term analysis, significantly enriched functional terms associated with the 43 genes regulated by both phytochromes under SD and LD included ‘transcriptional regulation’ and ‘photoperiodism’ (Additional file 1, Table S4). The 24 genes positively regulated by phytochromes (i.e. higher expression in WT than in *phy* mutants) included *FT1, FT2, FT3, PPD-B1, VRN1, FUL2* and *FUL3* (Figure 5, Additional file 1, Table S5). Although the effects were greater in LD, these results confirm that *PHYB* and *PHYC* also play a significant role in the activation of these genes in SD in the Kronos-PI background. These results are consistent with our qRT-PCR analysis (Figure 3). Other genes with the same expression profile as the previous group included a gene encoding a *CONSTANS*-like CCT-domain protein (TraesCS1A01G220300), and two homeologs encoding MYB-transcription factors with high similarity to *RADIALIS* (TraesCS6A01G273200 and TraesCS6B01G300600, Figure 5, Additional file 1, Table S5). One gene (TraesCS1A01G569000LC) was upregulated by *PHYB* in both SD and LD and by *PHYC* in SD, but was downregulated by *PHYC* in LDs (Additional file 4).

**Figure 5:**
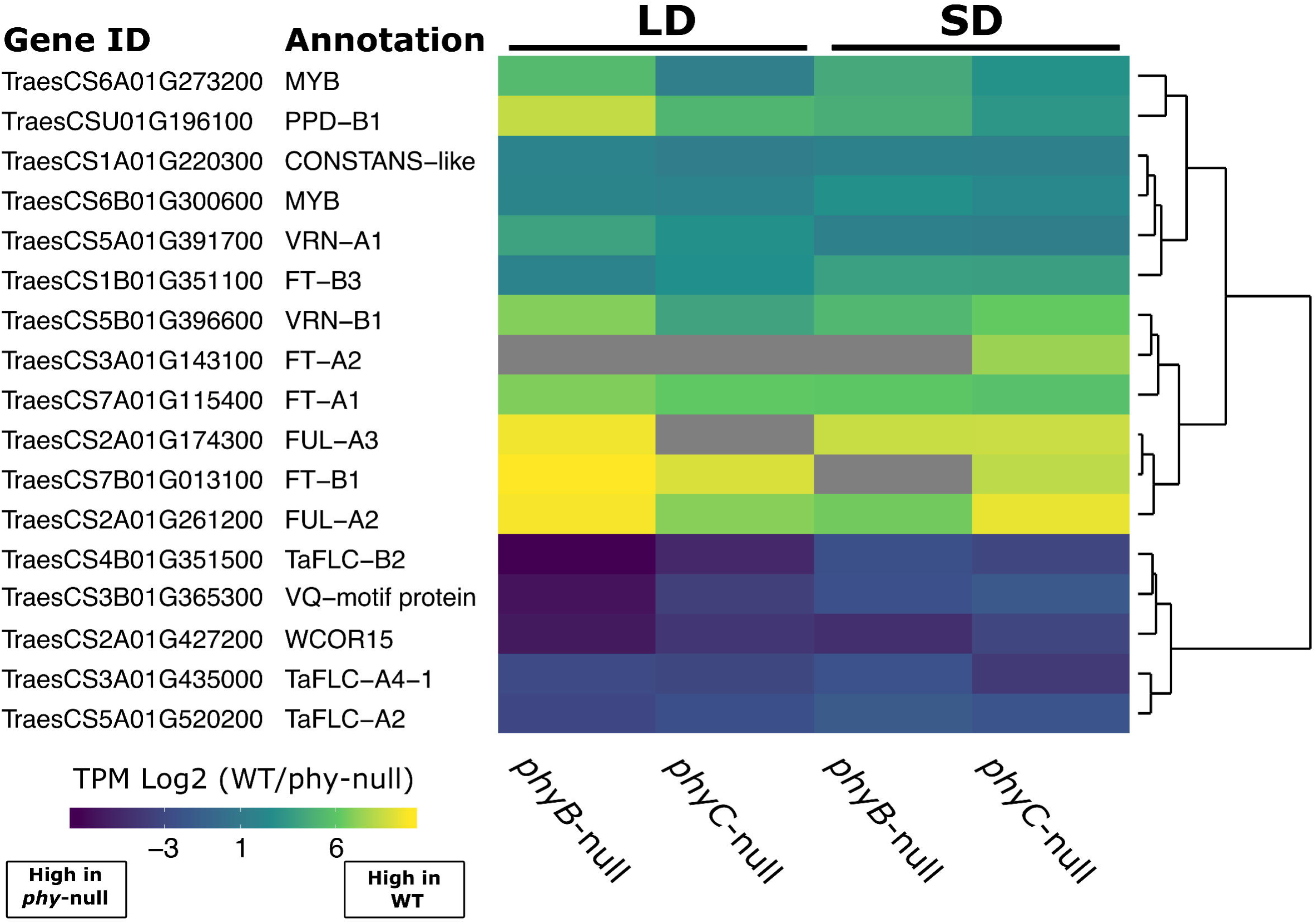
Heat map of relative expression changes of selected genes within the 43 DE genes regulated by both *PHYB* and *PHYC* in both SD and LD conditions. Expression values are presented as log2 TPM values of the fold-change between WT and each respective *phy* mutant. Gray color represents zero expression in the *phy* mutant.

Among the 17 genes that were negatively regulated by both *PHYB* and *PHYC* in both photoperiods were TraesCS3B01G365300, which encodes a member of the VQ motif protein family of transcriptional regulators, and three genes encoding members of the *FLC* clade of MADS-box TFs (Figure 5, Additional file 1, Table S5). Two of these genes encode homeologs of *FLC2*, which is orthologous to *OsMADS51*, a SD promoter of flowering in rice [33]. *FLC4* encodes the ortholog of *ODDSOC2*, which functions as a flowering repressor in *Brachypodium* and is induced by cold treatment in wheat [34]. Interestingly, TraesCS2A01G427200, which encodes *WCOR15*, a cold responsive gene, was strongly upregulated in both mutant lines, suggesting that phytochromes play an important role in suppressing the cold tolerance pathway in wheat under ambient temperature conditions (Figure 5, Additional file 1, Table S5). One gene (TraesCS2A01G019700LC) was downregulated by *PHYC* in both SD and LD and by *PHYC* in LD, but was upregulated by *PHYB* in LD conditions (Additional file 4).

A GO term analysis of the 589 genes regulated by both *PHYB* and *PHYC* only under SD, revealed enriched functional terms ‘protein phosphorylation’ and ‘homeostasis’ (Additional file 1, Table S4). Among the 367 genes positively regulated within this group, we detected nine WRKY transcription factors, both homeologs of a *RADIALIS*-like MYB-family transcription factor (TraesCS7A01G233300 and TraesCS7B01G131600) and TraesCS5B01G054800, which encodes a bHLH TF with similarity to the PIF subfamily (Figure 6a, Additional file 1, Table S6). We also found in this group *FT-A2*, *FT-A4* and *FLC-A1* (Figure 6a, Additional file 1, Table S6). Among the 213 genes that were negatively regulated by both phytochromes only under SD we identified members of the GATA, G2-like and B-box transcription factor families and TraesCS1A01G334400, which encodes the GA deactivating enzyme GA-2oxidase-A4 (Figure 6a, Additional file 1, Table S6). The upregulation of four members of the *CBF* family of cold-activated transcriptional regulators in both phytochrome mutants (Figure 6a, Additional file 1, Table S6), suggests a similar role to *WCOR15* in suppressing the cold tolerance pathway at ambient temperatures, but in this case restricted to SD conditions. Nineteen other genes were either positively regulated by *PHYB* and negatively regulated by *PHYC*, or *vice versa* (Additional file 4).

**Figure 6:**
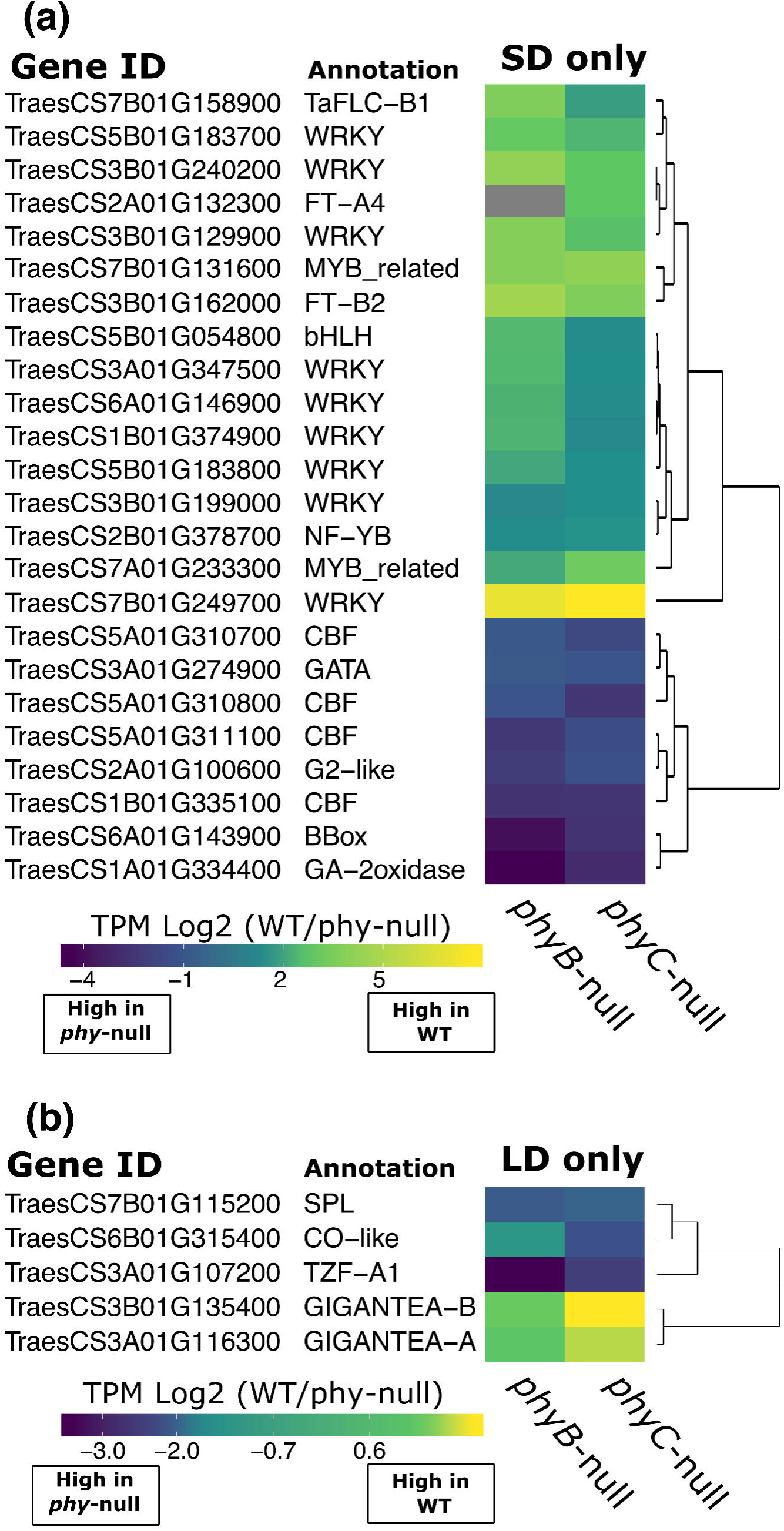
Heat map of relative expression changes of genes regulated by both *PHYB* and *PHYC* **(a)** specifically in SDs and **(b)** specifically in LDs. Expression values are presented as log2 TPM values of the fold-change between WT and each respective *phy* mutant. Gray color represents zero expression in the *phy* mutant.

We next studied the 46 genes regulated by both *PHYB* and *PHYC* specifically under LD conditions. Among the most significantly enriched functional terms associated with these genes were ‘shoot system development’, ‘long-day photoperiodism’ and ‘regulation of circadian rhythm’ (Additional file 1, Table S4). There were 27 genes positively regulated by both *PHYB* and *PHYC* in LD including both homeologs of *GIGANTEA*, suggesting this gene may play a role in the LD activation of flowering in wheat (Figure 6b, Additional file 1, Table S7). Among the 16 genes negatively regulated by both phytochromes was TraesCS6B01G315400, which encodes a *CONSTANS*-like protein, a member of the *SPL* family of transcription factors and *TANDEM ZINC FINGER1*, which, in Arabidopsis, interacts with PRR protein components of the circadian clock regulatory network [35] (Figure 6b, Additional file 1, Table S7). Three other genes were positively regulated by *PHYB* but negatively regulated by *PHYC* (Additional file 4).

### Effect of genotype on photoperiod regulated genes

Finally, we performed direct pairwise comparisons between SD and LD samples for each genotype (Additional file 5) to identify photoperiod-regulated genes (PRGs). There were a greater number of PRGs in both *phyB*-null (19,749) and *phyC*-null (13,740) mutants than in the wild-type (12,873, Additional file 1, Figure S4), suggesting that loss-of-function mutations in *phyB*-null and *phyC*-null were not sufficient to reduce the large effects on the wheat transcriptome generated by different photoperiods.

Although the different sampling points in LD (4 w) and SD (8 w) were selected so that WT genotypes were at similar developmental stages in both experiments, these results should be interpreted with caution because the effect of photoperiod is conflated with the effect of differences in chronological age. Both *phyB*-null and *phyC*-null mutants headed earlier under SD than under LD, so it is likely that the mutant lines were at different stages of development at the time of sampling. This particular sampling strategy likely contributed to the smaller number of PRGs in the WT genotypes relative to the *phyB*-null and *phyC*-null mutants.

We used this dataset to explore the expression profiles of 19 flowering time genes in different genotypes and photoperiods and the interaction between these factors (Figure 7).

**Figure 7:**
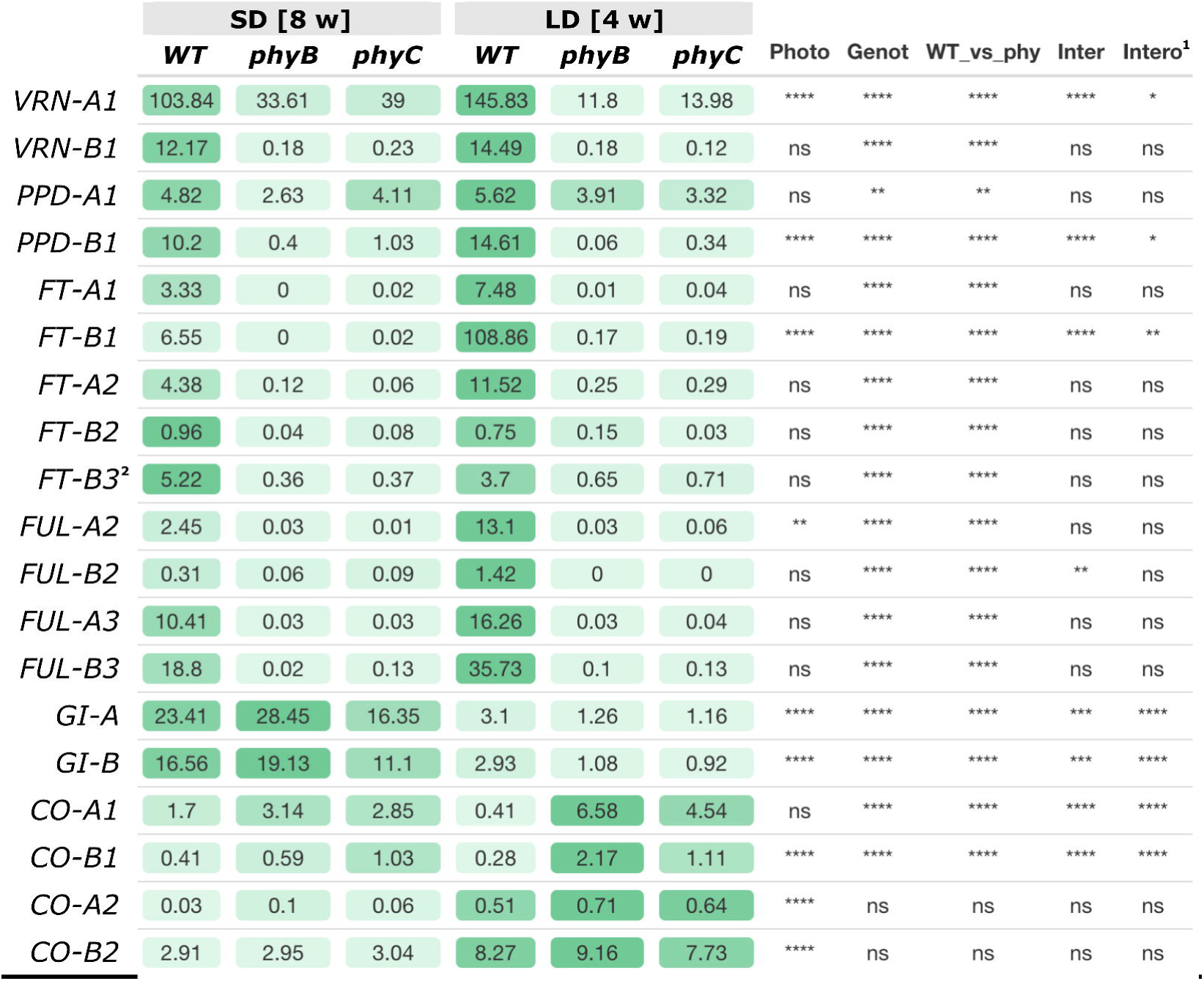
Photoperiod x Genotype factorial ANOVAs for transcripts per million (TPM) of 19 flowering time genes. Least square adjusted means of TPM (SD = 4 reps, LD = 8 reps) from the ANOVA are color coded so that higher transcript levels are indicated in darker shades of green (separately for each gene). WT *vs*. *phy* indicates an orthogonal contrast comparing the WT *versus* the two mutants. Data was transformed to provide normality of residuals. **** = *P* < 0.0001, *** = *P* < 0.001, ** = *P* < 0.01, * = *P* < 0.05, ns = not significant. ^1^ Since transformation affects the interpretation of the significance of the interactions, we also provide the significance of the interaction in the untransformed data. ^2^ *FT-A3* transcript levels were zero in all samples.

This analysis confirmed previous results showing that transcript levels of *VRN-A1, PPD-B1*, *FT-B1, FUL-A2*, *GI, CO-B1* and *CO2* are all significantly affected by photoperiod (Figure 7).

Notably, transcript levels of the photoperiod insensitive *Ppd-A1a* allele were not significantly affected by photoperiod in this dataset, whereas those of the *Ppd-B1b* showed a highly significant effect of photoperiod (*P* < 0.0001). Transcript levels of GI were more highly expressed in SD, whereas those of *CO1* and *CO2* were more highly expressed in LD (Figure 7).

The expression of most of these flowering time genes was also affected by the *phyB*-null and *phyC*-null mutations. Significant differences among the three genotypes were accompanied by significant differences between WT and the combined *phyB-* and *phyC*-null mutants, with the exception of *CO-A2* and *CO-B2*. The latter result is consistent with a previous study in which *CO1* was highly upregulated during the day in *phyC*-null mutants but *CO2* transcript levels were unaffected [28]. The *VRN1* paralogs (*FUL2* and *FUL3*) and the florigen-related genes (*FT1* and *FT2*) all share similar profiles, with higher transcript levels in the WT relative to the *phy-*null mutants and in LD relative to SD (Figure 7). *FT-A3* transcripts were not detected, whereas *FT-B3* transcript levels were higher in SD than in LD, consistent with the known role of this gene as a SD promoter of heading date [19, 36].

*VRN-A1*, *PPD-B1* and both homeologs of *GIGANTEA* were the only analyzed flowering promoting genes for which we observed significantly higher transcript levels in the *phy-*null mutants in SD than in LD. Based on this result, we speculate that these genes could contribute to the earlier flowering of the *phy-*null mutants in SD than in LD. Expression of these genes was significantly affected by photoperiod and genotype and all three showed significant genotype x photoperiod interactions (Figure 7). It is important to point out that the SD RNA-seq samples for the *phy-*null mutants were collected 70-78 days before heading, so they likely represent early stages of flowering induction. It would be interesting to study later time points closer to heading to see if genes that are induced by *VRN-A1*, such as *VRN-B1*, *FT1*, and *FT2* [32, 37], are upregulated earlier in SD than in LD.

### Light signaling and alternative splicing (AS) in wheat

In addition to the differences in transcript levels, we explored whether *PHYB* or *PHYC* regulate AS events in wheat using the replicate Multivariate Analysis of Transcript Splicing (rMATS) statistical method [38]. Our RNA-seq datasets show that both *PHYB* and *PHYC* regulate the expression of genes encoding components of the splicing machinery (Additional file 1, Table S8). For example, TraesCS2A01G122400, which encodes the large subunit of splicing factor U2AF, was downregulated in *phyB*-null mutants in both SD and LD conditions and TraesCS1B01G130200, which encodes an Arginine/serine-rich splicing factor, was upregulated in *phyC*-null mutants in both SD and LD (Additional file 1, Table S8). There were also several splicing-related genes regulated specifically under SD conditions. Three genes encoding splicing factor subunits were upregulated in both *phyB*-null and *phyC*-null mutants, while TraesCS1B01G125800, which encodes pre-mRNA-splicing factor cwc26, was significantly downregulated in both mutants in SD conditions (Additional file 1, Table S8).

To quantify the effect of these changes on AS in wheat, we first identified RNA-seq reads mapping to exon-intron junctions in annotated genes and calculated the frequency of AS events in five different categories (retained intron, skipped exon, alternative 5’ or 3’ splice sites and mutually exclusive exons). Comparing the frequency of each event between WT and mutant genotypes in different photoperiods, we found 5,175 AS events that were significantly regulated by either *PHYB* or *PHYC* (FDR *P-*adj < 0.05). The most commonly observed AS event was intron retention, followed by alternative 3’ splice sites (Figure 8a).

**Figure 8:**
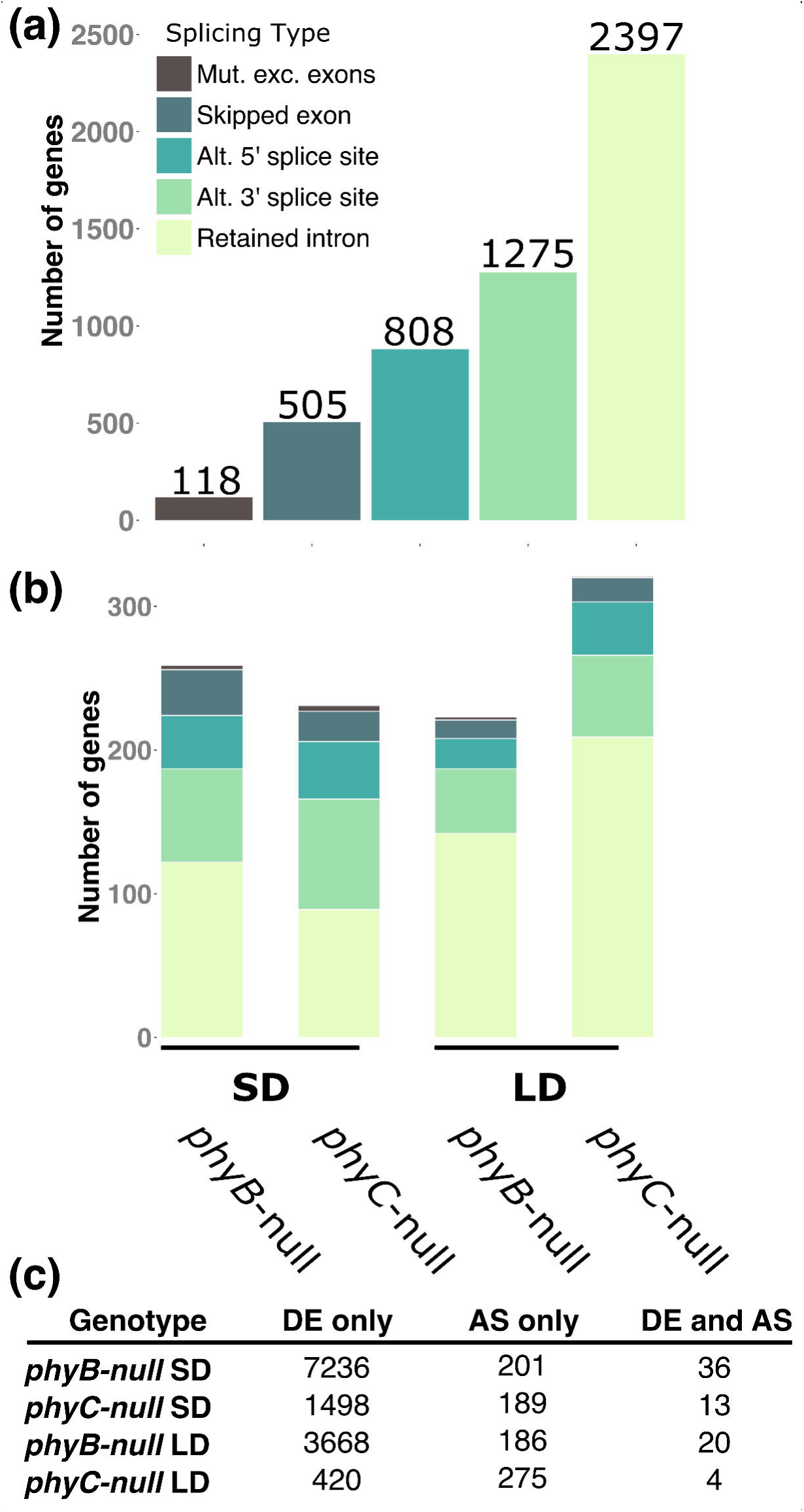
Phytochrome-mediated alternative splicing events in wheat **(a)** Number of AS events in each category among all RNA-seq data. **(b)** Number of genes differentially affected by AS events in WT and *phy*-null mutants in SD and LD RNA-seq experiments. **(c)** Overlap between DE genes and AS genes in pairwise comparisons between WT and *phy*-null mutants in SD and LD photoperiods.

To classify the events with potentially greater impact on gene function, we looked at the subset annotated genes that showed >30% variation in their isoform expression levels between genotypes. Among these genes, similar numbers were impacted by AS events in SD and LD (Figure 8b), although we found a slightly larger number of genes with retained intron events mediated by *PHYC* in LD conditions (Figure 8b). The total number of genes affected by at least one AS event with >30% variation in frequency between genotypes ranged from 202 (189 AS only +13 AS & DE) in WT vs *phyC*-null in SD conditions to 279 (275 AS only + 4 AS & DE) between the same genotypes in LD conditions (Figure 8c). These results indicate that in all pairwise comparisons the proportion of genes showing AS was much lower than the proportion of DE genes, and that only a small proportion of genes are both differentially expressed and subject to AS (Figure 8c).

Among all genes subject to AS, the functional term ‘RNA processing’ was significantly enriched in the GO term analysis as was ‘etioplast organization’ suggesting that some genes impacted by AS by phytochromes may be involved in chloroplast function (Additional file 1, Table S9). Full information of the individual genes impacted by different AS events are provided in Additional file 6. Specific examples include TraesCS2B01G140300, which encodes a CONSTANS-like protein. A retained intron event in this gene was significantly more abundant in WT plants than in *phyC*-null mutants in LDs. We detected a retained intron event in *FT-A10* in LD regulated by both *PHYB* and *PHYC*, and differentially expressed intron retention events in four different MADS-box genes in different pairwise comparisons (*TaFLC-A2*, *TaFLC-A5*, *TaSOC1-A5* and *TaAGL12-B1*).

## Discussion

### Phytochromes interact with *PPD1* in the regulation of wheat heading time

Across plant species, one well-characterized function of phytochromes is to regulate flowering time in response to changes in photoperiod. Previous studies have shown that Kronos *phyB*-null and *phyC*-null mutant plants grown under LD headed much later than the wild-type [28-30], and those results were confirmed here (Figure 1a-b). By contrast, loss-of-function mutations in the orthologous *PHY* genes in the SD grasses rice and sorghum result in earlier heading under LD [39-42]. Despite the opposite effect of the *phyB*-null and *phyC*-null mutants on heading time in SD-and LD-grasses, these two genes promote the expression of *PPD1/PRR37* in both groups of grasses. The difference between them seems to appear downstream of *PHYB* and *PHYC*, since under LD conditions *PPD1/PRR37* functions as a flowering repressor in rice [12] and sorghum [13] but as a flowering promoter in the temperate grasses [9-11].

One unexpected result from our study was that both *phyB-*null and *phyC-*null mutants headed earlier in SD than in LD, suggesting that these plants were behaving as if they were SD-plants. It is important to note that these experiments were all performed in the variety ‘Kronos’ which carries the PI (*Ppd-A1a*) allele. This *PPD1* allele has a deletion in its promoter region that encompass the binding site of the ELF3 protein repressor [5], resulting in ectopic expression of *PPD1* during the night [14], which is critical for the photoperiodic response as demonstrated in night-break experiments. Induction of *PPD1* in the middle of a 16 h night (SD) by a 15 m pulse of light accelerates heading time almost as much as a LD photoperiod [4]. In *Brachypodium*, it has been proposed that *PHYC* activation of *PPD1* is mediated by ELF3 [5], so the elimination of an ELF3 binding site in the *Ppd-A1a* allele in wheat may limit the transmission of the phytochrome signal to *PPD1*. This may explain the reduced differences between SD and LD observed in the *phyC-*null mutant compared to the WT (Figure 1a-b). It will be important to determine the effects of *phyB*-null and *phyC*-null mutations on heading date in the presence of the PS *Ppd-A1b* allele to test if the presence of the *Ppd-A1a* allele is necessary for the accelerated heading time in SD than in LD observed in the *phy* mutants. We have initiated the crosses to perform this experiment.

Although we still do not know the mechanism by which heading date is accelerated in SD relative to LD in the *phy-*null mutants, the expression studies provide some clues and point to a role of *PPD1*. This gene, together with its downstream target *VRN1*, both show a significant interaction between photoperiod and genotype, so that transcript levels are higher in LD in the WT, but higher in SD in both the *phyB*-null and *phyC*-null mutants (Figure 7). This result suggests that the modulation of *PPD1* expression and the differential regulation of *VRN1* may be part of the mechanism that promotes early flowering in SD in the *phy-*null mutants.

The acceleration of heading time under SD in the *phy-*null mutants has some similarities with SD-vernalization, but also some differences. In PS accessions of winter wheat and *Brachypodium*, an exposure to SD for 6-8 w at room temperature followed by LD replaces the need for vernalization to accelerate heading date [43-46], but this was not observed in PI wheat accessions [45]. By contrast, we observed SD acceleration in the Kronos-PI background in the presence of *phyB*-null or *phyC*-null mutations, which suggests that different regulatory mechanisms are likely involved in these two phenomena.

The temporal reversion in the order of activation of the *FT1* and *FT2* genes in the WT and *phy* mutants may also contribute to the earlier flowering of the *phy-*null mutants in SD. In the presence of the wild type phytochrome alleles, *FT1* is expressed to higher levels in Kronos-PI earlier in development than the *FT2* gene in SD (Figure 3) and LD [32]. However, in the *phyC-* null mutant under SD, *FT2* transcripts were upregulated earlier than *FT1.* By 17 weeks, when these plants were starting to head, *FT2* reached very high expression levels (>10-fold *ACTIN*) in both the *phyB-*null and *phyC-*null mutants. In growth chamber experiments under LD, under SD followed by LD conditions and in fall-planted field experiments, *ft2*-null alleles conferred only a small delay in heading date [32]. However, the role of *FT2* in the regulation of heading time under SD in a *phy-*null background requires additional studies.

Although *FT3* transcript levels were lower than other assayed genes, they were also upregulated earlier than *FT1* in both *phyB*-null and *phyC*-null mutants (Figure 3). In barley, overexpression of the orthologue *HvFT3* accelerates heading in LDs and promotes the transition of the shoot apical meristem from the vegetative to the reproductive stage in both SD and LD [47]. In *Brachypodium*, *BdFTL9*, a member of the *FT3* clade promotes flowering in SD conditions [46]. This protein forms a floral activation complex only in the absence of *BdFT1* (i.e. SD conditions), describing a possible mechanism by which diversity in the PEBP family can finely tune flowering time control according to photoperiod [48]. We identified several other members of the PEBP family that were upregulated in LD conditions (Additional file 5), for which it would be interesting to characterize their role in wheat heading date.

In addition to the PEBP genes, *GIGANTEA*, *VRN2/GHD7* and *CO* have been shown to play important roles in the photoperiod response in rice [49, 50]. *GIGANTEA* is a direct promoter of FT in Arabidopsis [51], and in rice *GIGANTEA* upregulates *CO* (Hd1) which activates the expression of *FT* [50, 52]. In this study, we show that wheat *GIGANTEA* was expressed at significantly higher levels under SD than under LD and was positively regulated by both *PHYB* and *PHYC* specifically under LD (Figure 7), suggesting that *GIGANTEA* may also play a role in the wheat photoperiod pathway. In rice, CO promotes flowering in SD in the presence of functional *GHD7/VRN2* or *PRR37/PPD1* alleles, and in LD in the *ghd7prr37* double mutant [49] providing an example of how mutations in these photoperiod genes can result in the reversion of the photoperiodic response. Both wheat *CO1* homeologs were highly upregulated in both *phy* mutants, whereas the *CO2* homeologs were not affected by the same mutations suggesting that these two paralogs are regulated differently by the phytochrome genes. Interestingly, *CO1* transcript levels were higher in SD in the WT and in LD in the *phy* mutants resulting in a strong interaction between genotype and photoperiod (Figure 7). We also identified TraesCS7A01G211300, that encodes the ortholog to *BdCONSTANS-Like 1* (Additional file 6). This gene is upregulated in LD in WT genotypes, but upregulated in SD in *phyB*-null and *phyC*-null mutants. Interestingly this gene was differentially expressed in the *Brachypodium elf3*-null mutant, suggesting that the TraesCS7A01G211300 and *BdCONSTANS-Like 1* orthologs may share similar regulatory mechanisms [5].

We are unable to draw conclusions on the role of the *VRN2* locus (duplicated genes *ZCCT1* and *ZCCT2*) because the functional *ZCCT-B2a* and *ZCCT-B2b* genes are not annotated in the reference genome used in our study, and the non-functional *ZCCT-A1* (TraesCS5A01G541300) and *ZCCT-A2* genes (TraesCS5A01G541200) were expressed at low levels (Additional file 5).

When analyzing the expression profiles of flowering time genes it is important to remember that the RNA-seq data represent a single time point during the day and during plant development of a very dynamic process of interactions among multiple flowering genes. Therefore, these expression profiles can change if analyzed at different times or developmental stages. Despite this limitation, the information generated for this single time point provided important insights on the complex networks that regulate wheat development in response to the phytochrome signals.

### Phytochromes affect plant architecture and vegetative development

In addition to flowering time, we found that mutations in *PHYB* and *PHYC* are associated with differences in vegetative development. In both SD and LD photoperiods, the leaves in the *phyB*-null and *phyC*-null mutants were longer and wider than in the wild-type suggesting a more extended or more robust growth (Figure 1e-f, [28, 31]). This is in contrast to the *phyC*-null mutant phenotype in *Brachypodium*. The first four leaves of *phyC*-null plants were shorter than WT in SDs, and not significantly different in length in LD conditions [30]. This discrepancy could be due to the stage of development, since in our study, we measured flag leaves and in *Brachypodium*, young leaves were studied.

The impact of these alleles on plant height was strikingly different between photoperiods. Whereas under LD conditions both *phyB*-null and *phyC*-null mutant lines were shorter than WT [31], in SDs both mutants were significantly taller (Figure 1g). Interestingly, although overall height of *phyB* and *phyC* were similar, the stem development in each mutant was different, with *phyB*-null mutants exhibiting a greater number of internodes (Figure 1g). There were several genes regulated by *PHYB* but not *PHYC* that may be associated with these phenotypic differences. In both SD and LD, transcript levels of *GA20ox-B2* and *GA20ox-B4*, which encode GA biosynthetic enzymes, were significantly higher in *phyB*-null mutants than either WT or *phyC*-null (Additional file 4).

Mutations in the phytochrome genes also affect plant morphology in the short-day grasses. Among rice plants grown in the field under non-inductive LD conditions, those with no functional phytochromes headed earlier, were shorter and had smaller panicles than sister lines with a functional *PHYC* in a *phyA phyB* background [41]. In the *phyA phyB* background, the *PhyC* gene also affected chlorophyll content, leaf angle and grain size, confirming the multiple pleiotropic effects of the phytochrome mutants in grasses. Sorghum plants carrying non-functional *phyB* alleles exhibit elongated hypoctyl growth in response to blue light [53] and increased stem elongation and internode number in inductive photoperiods [54]. Similarly, Arabidopsis, a LD-plant, exhibits elongated petioles in *phyB* mutants [55]. These vegetative phenotypes are characteristic of the shade avoidance response, which is mediated by *PHYB* in both SD and LD species. The similarities in phenotypes suggest that *PHYB* may play a role in shade avoidance pathways in wheat that is conserved in the other plant species described above. Some of the multiple genes differentially regulated in *PHYB* but not in *PHYC* may play a role in the shade avoidance response.

Our transcriptomic results are also consistent with previous studies that have established a link between phytochromes and the cold regulation pathway [56, 57]. In rice, *phyB*-null mutants exhibit improved cold tolerance [58] and in Arabidopsis, PIF3 binds to the promoters of *CBF* genes to suppress their expression [59]. We identified four *CBF* genes and two *COR* genes that were highly expressed in phytochrome null mutants in SDs (Figures 5 and 6). Transcript levels of four *COR* genes were significantly higher in SD than LD in WT and both *phy* mutants, but the differences were greater in the *phy* mutants. This demonstrates that in warm ambient temperatures, both *PHYB* and *PHYC* act to suppress the activation of the cold responsive pathway during the day. In wheat, a link between light quality and cold tolerance has previously been made [60] and suggests that the destabilization of phytochromes in response to FR light (commonly at higher levels in the dusk) or darkness improves the overall cold tolerance. It would be interesting to test the cold tolerance of the Kronos phytochrome mutants to confirm this activation at the physiological level.

### Conclusions

In wheat, *PHYB* and *PHYC* regulate vegetative development and flowering time with null-mutants for each gene showing a stronger delay in heading time under LD than under SD. We found that the flowering promoting genes *PPD-B1*, *VRN-A1* and *GIGANTEA* were more highly expressed in SD than LD in the *phy* mutants, and hypothesize that they may contribute to the earlier flowering time of these plants in SD than in LD. Our study provides insights into wheat light signaling pathways in inductive and non-inductive photoperiods and identifies a set of novel candidate genes to dissect the underlying developmental regulatory networks.

## Methods

### Plant materials, growth conditions and phenotypic measurements

All experiments were performed in the tetraploid *Triticum turgidum* L. *ssp.* durum Desf. variety ‘Kronos’ (genomes AABB). The *phyB-null* and *phyC-null* mutants were identified from EMS-mutagenized TILLING populations [61] and were described previously [28, 31]. Briefly, we combined null mutations in the A and B homeologs of each gene by marker assisted selection and performed two backcrosses to reduce background mutations. We self-pollinated the mutants for several generations, and used BC_2_F_4_ *phyB*-null and BC_2_F_5_ *phyC*-null mutants for the RNA-seq studies. Wild-type lines correspond to the same Kronos parent used in the backcross. All plants in this experiment carried the *Ppd-A1a* allele that confers reduced sensitivity to photoperiod [11]. All plants were grown in growth chambers (PGR15, Conviron, Manitoba, Canada) under SD conditions (8 h light/16 h dark) at 20 °C day/18 °C night temperatures and a light intensity of ~260 µM m^−2^ s^−1^. All chambers used similar halide light configurations and were located in the same room.

Heading time was recorded as the number of days after sowing when half of the spike emerged from the boot (Zadoks 55 [62]) using five biological replications (n) per genotype. At maturity we measured total height and individual internode length (n=4), total tiller number, leaf number, flag leaf width and length (n=6). We compared SD data for heading time with previously published heading data of the same mutants grown in the same growth chamber configuration under LD [31].

### qRT-PCR assays

Beginning when plants were two-weeks old, we collected tissue from the last fully expanded leaf in liquid nitrogen at three-week intervals until 17 weeks after sowing to cover most of the developmental stages in the mutants. We collected four biological replicateof all three genotypes (wild-type, *phyB*-null and *phyC*-null) at each time point. We extracted RNA using the Spectrum™ Plant Total RNA kit (Sigma-Aldrich, St. Louis, MO) following the manufacturer’s instructions. cDNAs were synthesized from 1 µg of total RNA using the High Capacity Reverse Transcription Kit (Applied Biosystems) and quantitative RT-PCR was performed in a 7500 Fast Real-Time PCR system (Applied Biosystems, Foster City, CA) using SYBR Green. Primers for the target genes *PPD1* [28], *FT1* [15], *FT2* [16], *FT-A3*, *FT-B3* [63], *VRN1* [15], and the control gene *ACTIN* [31] were described previously. Expression data are presented as fold-*ACTIN* levels (molecules of target gene/molecules of *ACTIN*).

### RNA-seq library construction and sequencing

The individual plants used for the RNA-seq experiment were the same plants used for the qRT-PCR and phenotypic studies. For the SD RNA-seq experiment, we extracted RNA samples from eight-week-old plants. At this stage, the apices of the wild-type plants were at an early stage of spike development (Waddington stage 3 [64]) and the apices of both *phyB*-null and *phyC*-null plants were still in the vegetative stage (Waddington stage 1 [64]). Data from the LD RNA-seq experiment was previously described [31] and was generated from RNA extracted from the fully-extended third leaf of four-week-old plants, when the apices of wild-type plants were at the same developmental stage as in eight-week-old SD-plants. We assembled RNA-sequencing libraries using the TruSeq RNA Sample Preparation kit v2 (Illumina, San Diego, CA), according to the manufacturer’s instructions. Library quality was determined using a high-sensitivity DNA chip run on a 2100 Bioanalyzer (Agilent Technologies, Santa Clara, CA). Libraries were barcoded to allow multiplexing and were sequenced using the 100 bp single read module across two lanes (two biological replicates of each genotype (= six libraries) per lane on a HiSeq4000 sequencer at the UC Davis Genome Center.

### RNA-seq data processing

Raw reads were processed using a pipeline incorporating “*Scythe*” (https://github.com/vsbuffalo) to remove Illumina adapter contamination (default options) and “*Scythe*” (https://github.com/najoshi/sickle) to remove low-quality reads (With options -t sanger -q 25 -l 50). Processed reads were mapped to the IWGSC RefSeq v1.0 genome assembly [65], using GSNAPl [66]. We used parameters -m 4 -n 1 -A sam -N 1 -t 24 for the 100 bp single end read SD data, and parameters -m 2 -n 1 -A sam -N 1 -t 24 for the 50 bp single end read LD data, to generate Sequence Alignment/Map (SAM) files for each sample. We used high and low confidence gene models from IWGSC Refseq v1.0 gene models. To provide additional context to gene function, we performed a BLASTP search using each annotated gene as a query against the NCBI NR database of proteins. We also added additional annotation information for genes encoding members of different transcription factor families [67], MIKC subclass members of the MADS-box gene family [68] and of the *FT-like* gene family [63]. Full information of the annotations associated with each differentially expressed gene are provided in Additional file 2.

Raw count values were generated using htseq-count (https://github.com/simon-anders/htseq) on each of the resulting SAM files, using the options -m union --stranded=no -a 40 -t gene -i ID. These mapping parameters ensured that reads with an alignment quality lower than 40 were discarded, so that only counts from uniquely mapped reads were considered for gene expression analyses. Genes that showed no raw count values greater than or equal to three in any replicate of any of the three genotypes were discarded, leaving 72,108 genes with a level of expression above our threshold. The raw counts for these remaining genes were normalized using DESeq2. After normalization, we applied the statistical tests implemented in both DESeq2 and edgeR to classify differentially expressed genes in pairwise comparisons. The P-values generated by both analyses were adjusted for FDR, using the procedure of Benjamini and Hochberg [69] and we selected a stringent cutoff of adjusted *P ≤* 0.01 for significance for both tests within each experimental replication. For LD data, two experimental replicates were analyzed separately and only genes that were significant in both comparisons (described as “high-confidence” DE genes in our earlier study), were included in this analysis.

### Alternative splicing

Alternative splicing events were characterized with rMATS v4.0.1 [38]. A GTF annotation file was created for both SD and LD datasets using Stringtie [70]. Inputs for this file were the sorted BAM files generated during RNA-seq mapping and high and low confidence gene annotations from IWGSC RefSeq v1.1 to specify exon-intron boundaries. Genome indices used by rMATS were created from the IWGSC RefSeq v1.0 assembly using STAR (parameter --runMode genomeGenerate) [71]. Fastq files for each sample were trimmed to 100bp and 50bp for SD and LD datasets, respectively, using a custom perl script. rMATS was run twice on each dataset, comparing WT with *phyB*-null and WT with *phyC*-null samples in both SD and LD datasets, using their respective GTF annotation files [65]. The inclusion level difference for each alternative splicing event was calculated from the number of reads for each replicate that map to a possible inclusion event, normalized by the length of those possible events. The value for each type of event represents the pairwise comparisons of the mean value from four replicates of wild-type and the respective *phy*-null genotype. Positive inclusion level differences indicate more reads mapped to an AS event in wild type than in the *phy*-null sample and vice versa. An initial 0.01% splicing difference and FDR < 0.05 filter was used to determine significant alternative splicing events categorized into retained introns, skipped exons, alternative 5’ splice sites, alternative 3’ splice sites, and mutually exclusive exons. A more stringent cutoff of 30% inclusion level difference was used to analyze a subset of these events in greater detail.

### Functional annotation

We identified the longest transcribed contig mapping to each genomic locus and performed a BLASTX against the nr protein database (nr.28, Apr 24, 2015 release, NCBI) and a BLASTP using the translated ORF against the Pfam database version 27.0 with InterProScan version 5.13 to identify conserved protein domains. The output was used to infer GO terms associated with each genomic locus using BLAST2GO version 2.6.5 and we used the ‘R’ package TopGO version 2.14.0 to perform an enrichment analysis among the differentially regulated gene sets. “Biological Process” terms were obtained and significance values for enrichment were calculated using ‘classic’ Fishers’ exact test, as implemented in TopGO.

## List of abbreviations

AS: Alternative Splicing
BAM: Binary Alignment Map
bHLH: basic Helix Loop Helix
COR: Cold Responsive
DE: Differentially Expressed
EMS: Ethyl-Methane Sulfonate
FDR: False Discovery Rate
FR: Far Red
GO: Gene Ontology
IWGSC: International Wheat Genome Sequencing Consortium
LD: Long Day
MDS: Multi-Dimensional Scaling
PEBP: Phosphatidylethanolamine-Binding Proteins
PHY: Phytochrome
PIF: Phytochrome Interacting Factor
PRG: Photoperiod Regulated Gene
PRR: Pseudo-Response Regulator
PI: Photoperiod Insensitive
PS: Photoperiod Sensitive
qRT-PCR: quantitative Reverse Transcriptase Polymerase Chain Reaction
R: Red
rMATS: replicate Multivariate Analysis of Transcript Splicing
SAM: Sequence Alignment Map
SEM: Standard Error of the Mean
TILLING: Targeted Induced Local Lesions IN Genomes
TPM: Transcripts per Million
WT: Wild-type

## Declarations

### Availability of data and materials

RNA-seq reads and raw count data is available at NCBI GEO (https://www.ncbi.nlm.nih.gov/geo/) under the accession number GSE141000. Previously published raw data from RNA-seq studies is available under accession number GSE79049. Genetic materials from this study are available by request.

### Author contributions

NK developed plant materials, performed all phenotypic analyses and molecular experiments, performed data analysis and contributed to writing the manuscript. CVG, JH, AA, HB performed expression data analysis. JD contributed to the initial coordination of the project, to data analyses and to the writing of the manuscript. SP performed data analysis and wrote the manuscript. All authors read and approved the final manuscript.

## Additional files

Additional file 1: Figures S1-S4, Tables S1-S9 (.pdf).

Additional file 2: RNA-seq data for all samples from SD photoperiods (.xls).

Additional file 3: RNA-seq data for all samples from LD photoperiods (.xls).

Additional file 4: RNA-seq data and annotations of genes regulated by *PHYB* or *PHYC* under SD or LD, divided into mutually exclusive categories (.xls).

Additional file 5: RNA-seq data comparing SD and LD within genotypes (.xls).

Additional file 6: Alternative splicing data from all pairwise comparisons (.xls).

