## Additional file 1 for "Effect of *phyB* and *phyC* loss-of-function mutations on the wheat transcriptome under short and long day photoperiods"

Additional file 1. Tables and figures in this additional file:

**Figure S1:** Heading date of *phyB* and *phyC* mutants in SD photoperiods. For each phytochrome gene, data is presented for plants carrying non-functional copies of only the A homeolog (-A), only the B homoeologue (-B) or both A and B alleles combined (null). Experiments were performed in the tetraploid variety 'Kronos' (Genomes AABB).

**Figure S2:** Representative plants of WT, *phyB*-null and *phyC*-null genotypes at eight-weeks of age. SD RNA-seq data was generated from leaf tissue harvested at this stage.

**Figure S3:** MDS plot for RNA-seq samples under LD conditions. Two experimental replicates consisting of four biological replicates of each genotype were processed.

**Figure S4:** Number of DE genes between SD and LD samples of each genotype. SD samples were harvested at 8 weeks of age and consisted of 4 biological replicates. LD samples were harvested at 4 weeks of age and consisted of 8 biological replicates.

**Table S1:** Summary of RNA-seq reads and mapping rates from SD samples.

**Table S2 –** Top 10 significant enriched GO terms among genes differentially regulated by *PHYB*, by *PHYC*, and by both *PHYB* and *PHYC* in SD conditions.

**Table S3:** Summary of RNA-seq reads and mapping rates from LD samples.

**Table S4 –** Top 10 significant enriched GO terms among genes differentially regulated by both *PHYB* and *PHYC* under different photoperiod conditions.

**Table S5**: Selected genes regulated by both *PHYB* and *PHYC* in both SD and LD photoperiods. Fold change between WT and respective *phy*-null mutants are presented. The values for LD are the mean fold-change in expression between both experimental replicates. a - *FT-A1* and *FT-B1* expression was zero in the *phy*-null mutants in these comparisons.

**Table S6**: Selected genes regulated by both *PHYB* and *PHYC* in SD photoperiods but not LD photoperiods. Fold change between WT and respective *phy*-null mutants are presented as the mean fold-change between both experimental replicates. a - *FT-A4* expression was zero in the *phy*-null mutants in these comparisons.

**Table S7**: Selected genes regulated by both *PHYB* and *PHYC* in LD photoperiods but not SD photoperiods. Fold change between WT and respective *phy*-null mutants are presented as the mean fold-change between both experimental replicates.

**Table S8:** *PHYB*- and *PHYC*-regulated splicing-related genes. Mean TPM and fold-change for each genotype was calculated from four biological replicates in SD samples, and eight biological replicates for LD samples.

**Table S9:** Top 40 significant enriched GO terms among genes impacted by any AS event by either *PHYB* or *PHYC* under SD or LD conditions.

**Figure S1:** Heading date of *phyB* and *phyC* mutants in SD photoperiods. For each phytochrome gene, data is presented for plants carrying non-functional copies of only the A homeolog (-A), only the B homoeologue (-B) or both A and B alleles combined (null). Experiments were performed in the tetraploid variety 'Kronos' (Genomes AABB).

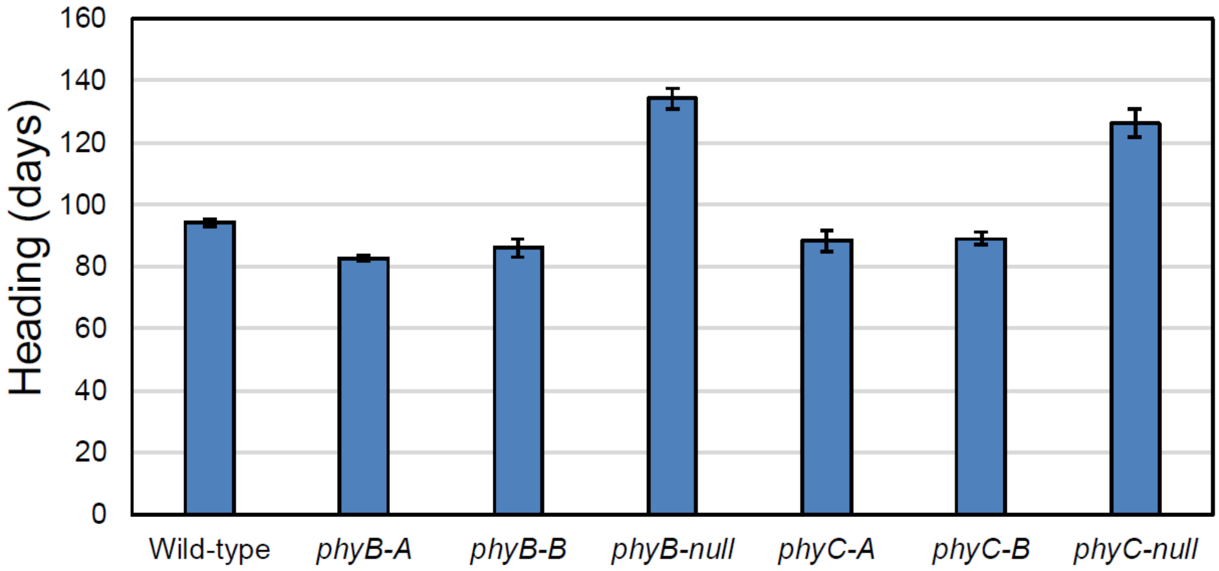

**Figure S2:** Representative plants of WT, *phyB*-null and *phyC*-null genotypes at eight-weeks of age. SD RNA-seq data was generated from leaf tissue harvested at this stage.

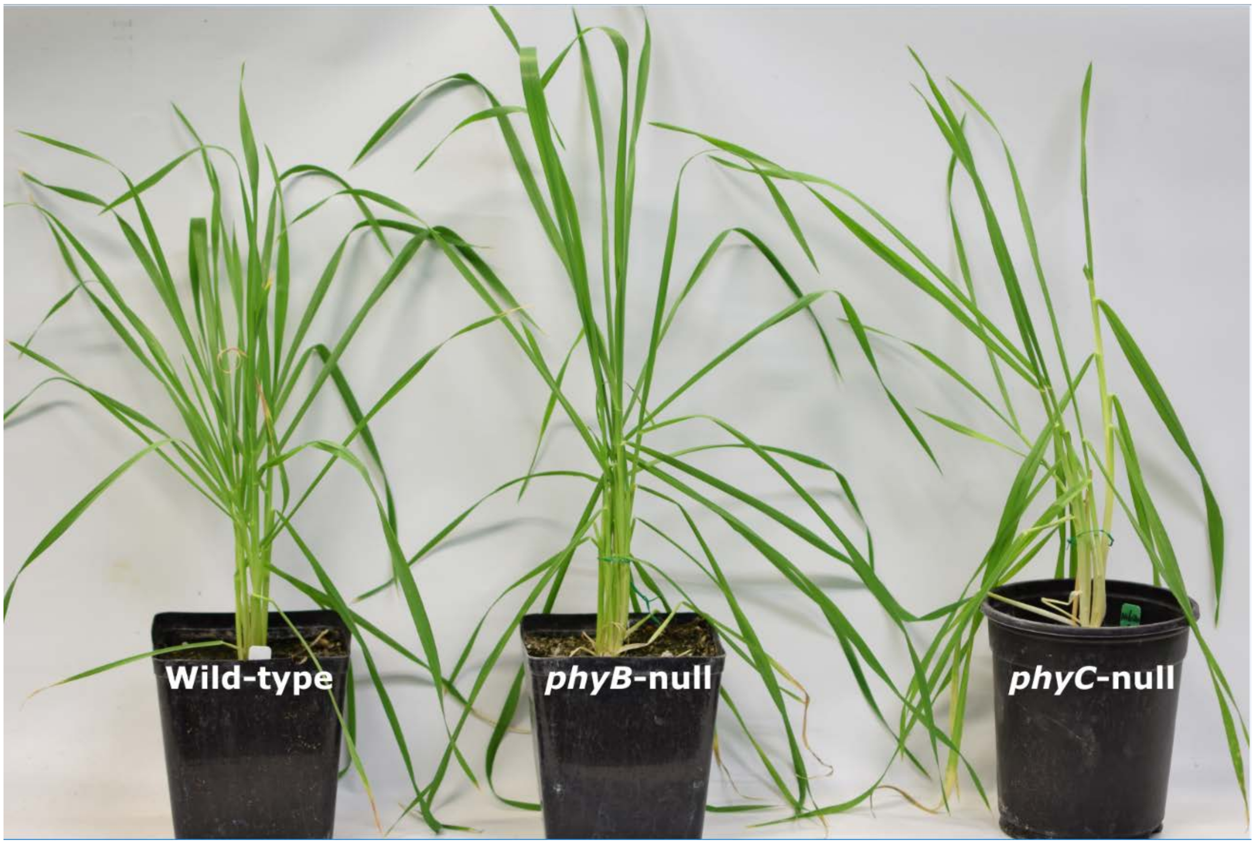

**Figure S3:** MDS plot for RNA-seq samples under LD conditions. Two experimental replicates consisting of four biological replicates of each genotype were processed.

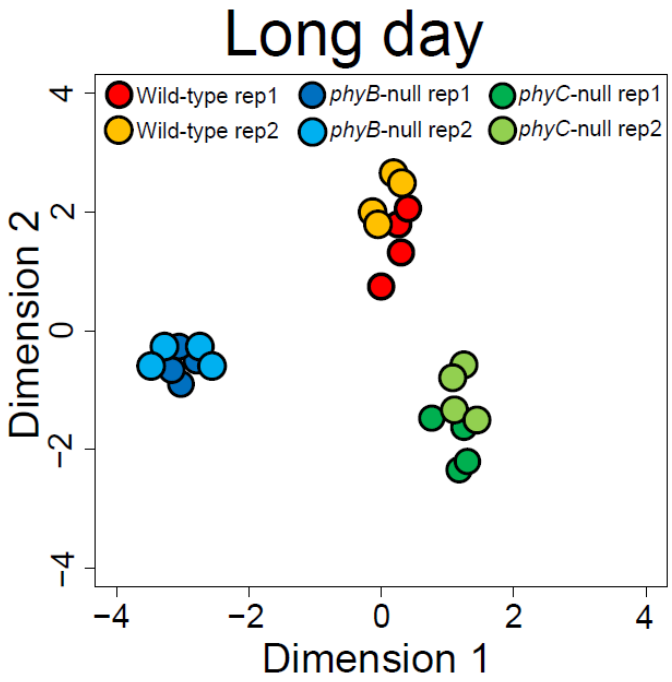

**Figure S4:** Number of DE genes between SD and LD samples of each genotype. SD samples were harvested at 8 weeks of age and consisted of 4 biological replicates. LD samples were harvested at 4 weeks of age and consisted of 8 biological replicates.

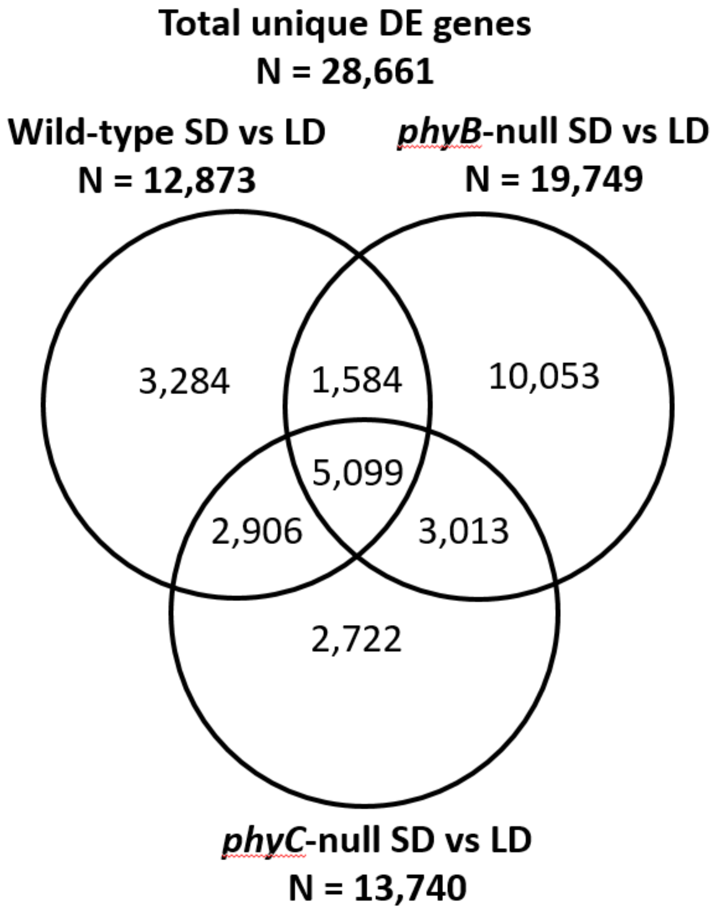

**Table S1:** Summary of RNA-seq reads and mapping rates from SD samples.

| Genotype | Biological replicate | Total raw reads | Reads discarded | | Total trimmed reads | Uniquely mapped reads | |
|---|---|---|---|---|---|---|---|
| | | | N | % | | N | % |
| **Wild type** | 1 | 58,502,128 | 1,013,691 | 1.7 | 57,488,437 | 39,607,623 | 68.9 |
| | 2 | 58,830,113 | 1,108,319 | 1.9 | 57,721,794 | 40,219,742 | 69.7 |
| | 3 | 67,577,803 | 1,139,114 | 1.7 | 66,438,689 | 45,702,198 | 68.8 |
| | 4 | 65,430,265 | 1,191,428 | 1.8 | 64,238,837 | 44,160,635 | 68.7 |
| ***phyB*-null** | 1 | 54,007,718 | 837,760 | 1.6 | 53,169,958 | 37,368,473 | 70.3 |
| | 2 | 63,069,027 | 1,079,413 | 1.7 | 61,989,614 | 43,188,697 | 69.7 |
| | 3 | 64,820,390 | 1,047,399 | 1.6 | 63,772,991 | 44,409,244 | 69.6 |
| | 4 | 66,273,908 | 1,174,901 | 1.8 | 65,099,007 | 45,971,041 | 70.6 |
| ***phyC*-null** | 1 | 66,985,324 | 1,317,576 | 2.0 | 65,667,748 | 45,956,376 | 70.0 |
| | 2 | 66,376,725 | 1,289,943 | 1.9 | 65,086,782 | 45,148,294 | 69.4 |
| | 3 | 86,487,404 | 1,503,903 | 1.7 | 84,983,501 | 58,948,262 | 69.4 |
| | 4 | 72,794,612 | 1,384,225 | 1.9 | 71,410,387 | 49,720,482 | 69.6 |
| **Mean** | | 65,929,618.08 | 1,173,972.67 | 1.8 | 64,755,645 | 45,033,422 | 69.7 |

**Table S2 –** Top 10 significant enriched GO terms among genes differentially regulated by *PHYB*, by *PHYC*, and by both *PHYB* and *PHYC* in SD conditions.

| ***PHYB*-regulated** | | | | | |
|---|---|---|---|---|---|
| **GO ID** | **Term** | **Annotated** | **Significant** | **Expected** | ***P*** |
| GO:0055114 | Oxidation-reduction process | 4,000 | 540 | 396.1 | 5.30E-15 |
| GO:0006468 | Protein phosphorylation | 3,908 | 529 | 387.0 | 6.60E-15 |
| GO:0044710 | Single-organism metabolic process | 9,737 | 1,161 | 964.1 | 8.50E-15 |
| GO:1903825 | Organic acid transmembrane trans. | 244 | 64 | 24.2 | 2.30E-13 |
| GO:1905039 | Carboxylic acid transmembrane trans. | 237 | 61 | 23.5 | 2.00E-12 |
| GO:0015849 | Organic acid transport | 324 | 75 | 32.1 | 2.10E-12 |
| GO:0009812 | Flavonoid metabolic process | 489 | 98 | 48.4 | 9.80E-12 |
| GO:0046942 | Carboxylic acid transport | 317 | 72 | 31.4 | 1.50E-11 |
| GO:0016310 | Phosphorylation | 5,363 | 669 | 531.0 | 1.70E-11 |
| GO:0044699 | Single-organism process | 17,160 | 1,886 | 1699.1 | 2.30E-11 |
| ***PHYC*-regulated** | | | | | |
| GO:0006952 | Defense response | 1,209 | 43 | 19.0 | 7.70E-07 |
| GO:0042592 | Homeostatic process | 1,321 | 43 | 20.8 | 7.20E-06 |
| GO:0006468 | Protein phosphorylation | 3,908 | 94 | 61.6 | 2.10E-05 |
| GO:0016310 | Phosphorylation | 5,363 | 121 | 84.5 | 2.20E-05 |
| GO:0016045 | Detection of bacterium | 5 | 3 | 0.1 | 3.80E-05 |
| GO:0098543 | Detection of other organism | 5 | 3 | 0.1 | 3.80E-05 |
| GO:0098581 | Detection of external biotic stimulus | 8 | 3 | 0.1 | 0.00021 |
| GO:0065008 | Regulation of biological quality | 2,033 | 53 | 32.0 | 0.00025 |
| GO:0048878 | Chemical homeostasis | 621 | 22 | 9.8 | 0.00041 |
| GO:*0006879* | Cellular iron ion homeostasis | 42 | 5 | 0.7 | 0.0005 |
| ***PHYB*/*PHYC*-regulated** | | | | | |
| GO:0006952 | Defense response | 1,209 | 30 | 11.1 | 9.10E-07 |
| GO:0006468 | Protein phosphorylation | 3,908 | 62 | 35.7 | 1.10E-05 |
| GO:0006879 | Cellular iron ion homeostasis | 42 | 5 | 0.4 | 4.00E-05 |
| GO:0016310 | Phosphorylation | 5,363 | 76 | 49.0 | 5.10E-05 |
| GO:0055072 | Iron ion homeostasis | 92 | 6 | 0.8 | 0.00021 |
| GO:0050896 | Response to stimulus | 7,059 | 90 | 64.5 | 0.0004 |
| GO:0042592 | Homeostatic process | 1,321 | 25 | 12.1 | 0.00054 |
| GO:0006950 | Response to stress | 3,927 | 54 | 35.9 | 0.00153 |
| GO:0048872 | Homeostasis of number of cells | 7 | 2 | 0.1 | 0.0017 |
| GO:0009812 | Flavonoid metabolic process | 489 | 12 | 4.5 | 0.00199 |

**Table S3:** Summary of RNA-seq reads and mapping rates from LD samples.

| Genotype | Exp. Rep. | Bio. Rep. | Raw reads | Reads discarded | | Trimmed reads | Uniquely mapped reads | |
|---|---|---|---|---|---|---|---|---|
| | | | | N | % | | N | % |
| **PHYB-WT** | **1** | 1 | 38,193,836 | 2,291,251 | 6.0 | 35,902,585 | 18,977,319 | 52.9 |
| | | 2 | 37,228,077 | 2,236,373 | 6.0 | 34,991,704 | 18,615,718 | 53.2 |
| | | 3 | 50,793,083 | 2,966,158 | 5.8 | 47,826,925 | 25,410,013 | 53.1 |
| | | 4 | 44,301,543 | 2,715,483 | 6.1 | 41,586,060 | 21,258,178 | 51.1 |
| | **2** | 1 | 53,475,278 | 3,983,069 | 7.5 | 49,492,209 | 26,092,448 | 52.7 |
| | | 2 | 54,275,607 | 3,972,114 | 7.3 | 50,303,493 | 26,657,660 | 53.0 |
| | | 3 | 57,095,077 | 4,313,269 | 7.6 | 52,781,808 | 27,415,963 | 51.9 |
| | | 4 | 59,581,947 | 4,279,495 | 7.1 | 55,302,452 | 29,303,043 | 53.0 |
| ***phyB-null*** | **1** | 1 | 43,374,525 | 2,540,401 | 5.9 | 40,834,124 | 21,739,116 | 53.2 |
| | | 2 | 47,201,620 | 2,731,653 | 5.8 | 44,469,967 | 23,537,045 | 52.9 |
| | | 3 | 81,109,422 | 4,903,617 | 6.0 | 76,205,805 | 40,552,227 | 53.2 |
| | | 4 | 47,250,183 | 2,769,089 | 5.9 | 44,481,094 | 23,788,301 | 53.5 |
| | **2** | 1 | 47,550,092 | 3,517,065 | 7.4 | 44,033,027 | 23,533,579 | 53.5 |
| | | 2 | 50,880,434 | 3,783,852 | 7.4 | 47,096,582 | 25,126,305 | 53.4 |
| | | 3 | 56,371,576 | 4,144,097 | 7.4 | 52,227,479 | 27,938,705 | 53.5 |
| | | 4 | 51,113,770 | 3,679,790 | 7.2 | 47,433,980 | 25,437,698 | 53.6 |
| **PHYC-WT** | **1** | 1 | 40,263,724 | 4,609,655 | 11.5 | 35,654,069 | 18,918,318 | 53.1 |
| | | 2 | 40,409,065 | 4,363,202 | 10.8 | 36,045,863 | 20,341,146 | 56.4 |
| | | 3 | 42,605,342 | 4,775,194 | 11.2 | 37,830,148 | 20,222,651 | 53.5 |
| | | 4 | 39,796,668 | 4,597,431 | 11.6 | 35,199,237 | 18,893,499 | 53.7 |
| | **2** | 1 | 37,994,979 | 3,339,800 | 8.8 | 34,655,179 | 18,184,337 | 52.5 |
| | | 2 | 43,262,009 | 3,678,319 | 8.5 | 39,583,690 | 21,115,165 | 53.3 |
| | | 3 | 41,619,486 | 3,613,203 | 8.7 | 38,006,283 | 20,202,760 | 53.2 |
| | | 4 | 31,887,538 | 2,770,104 | 8.7 | 29,117,434 | 15,386,865 | 52.8 |
| ***phyC-null*** | **1** | 1 | 44,904,496 | 4,989,736 | 11.1 | 39,914,760 | 21,705,064 | 54.4 |
| | | 2 | 46,776,974 | 5,163,917 | 11.0 | 41,613,057 | 22,393,932 | 53.8 |
| | | 3 | 43,960,996 | 4,795,476 | 10.9 | 39,165,520 | 21,053,322 | 53.8 |
| | | 4 | 42,873,364 | 4,595,730 | 10.7 | 38,277,634 | 20,718,521 | 54.1 |
| | **2** | 1 | 35,291,647 | 2,998,278 | 8.5 | 32,293,369 | 17,279,830 | 53.5 |
| | | 2 | 36,712,107 | 3,016,466 | 8.2 | 33,695,641 | 17,747,367 | 52.7 |
| | | 3 | 45,656,274 | 3,815,739 | 8.4 | 41,840,535 | 22,551,769 | 53.9 |
| | | 4 | 38,922,327 | 3,391,092 | 8.7 | 35,531,235 | 13,034,303 | 36.7 |
| Mean | | | 46,022,908 | 3,729,379 | 8.2 | 42,293,530 | 22,347,880 | 52.8 |

**Table S4 –** Top 10 significant enriched GO terms among genes differentially regulated by both *PHYB* and *PHYC* under different photoperiod conditions.

| ***PHYB-PHYC* regulated – SD specific** | | | | | |
|---|---|---|---|---|---|
| **GO ID** | **Term** | **Annotated** | **Significant** | **Expected** | ***P*** |
| GO:0006468 | Protein phosphorylation | 5,201 | 88 | 41.65 | 5.10E-12 |
| GO:0016310 | Phosphorylation | 7,036 | 105 | 56.35 | 4.60E-11 |
| GO:0019725 | Cellular homeostasis | 845 | 23 | 6.77 | 4.50E-07 |
| GO:0006952 | Defense response | 1,563 | 33 | 12.52 | 5.10E-07 |
| GO:0042592 | Homeostatic process | 1,597 | 33 | 12.79 | 8.10E-07 |
| GO:0006796 | Phosphate-containing compound metabolic process | 9,005 | 109 | 72.12 | 2.00E-06 |
| GO:0006793 | Phosphorus metabolic process | 9,117 | 109 | 73.02 | 3.60E-06 |
| GO:0045454 | Cell redox homeostasis | 356 | 13 | 2.85 | 7.20E-06 |
| GO:0006464 | Cellular protein modification process | 8,493 | 99 | 68.02 | 3.60E-05 |
| GO:0036211 | Protein modification process | 8,493 | 99 | 68.02 | 3.60E-05 |
| ***PHYB-PHYC* regulated – LD specific** | | | | | |
| GO:0048586 | Regulation of LD photoperiodism, flowering | 30 | 2 | 0.02 | 0.00015 |
| GO:0042752 | Regulation of circadian rhythm | 31 | 2 | 0.02 | 0.00016 |
| GO:2000028 | Regulation of photoperiodism, flowering | 41 | 2 | 0.02 | 0.00029 |
| GO:2000243 | Positive regulation of reproductive process | 42 | 2 | 0.03 | 0.0003 |
| GO:0048574 | LD photoperiodism, flowering | 43 | 2 | 0.03 | 0.00032 |
| GO:0005982 | Starch metabolic process | 224 | 3 | 0.14 | 0.00034 |
| GO:0048571 | LD photoperiodism | 45 | 2 | 0.03 | 0.00035 |
| GO:0010218 | Response to far red light | 52 | 2 | 0.03 | 0.00046 |
| GO:0007623 | Circadian rhythm | 53 | 2 | 0.03 | 0.00048 |
| GO:0048511 | Rhythmic process | 57 | 2 | 0.03 | 0.00056 |
| ***PHYB-PHYC* regulated – SD and LD** | | | | | |
| GO:0050789 | Regulation of biological process | 10,086 | 17 | 5.9 | 3.20E-06 |
| GO:0009889 | Regulation of biosynthetic process | 5,573 | 12 | 3.26 | 2.40E-05 |
| GO:0065007 | Biological regulation | 11,745 | 17 | 6.88 | 2.90E-05 |
| GO:0006355 | Regulation of transcription, DNA-templated | 5,066 | 11 | 2.97 | 5.80E-05 |
| GO:0080090 | Regulation of primary metabolic process | 6,135 | 12 | 3.59 | 6.40E-05 |
| GO:1903506 | Regulation of nucleic acid-templated transcription | 5,113 | 11 | 2.99 | 6.40E-05 |
| GO:2001141 | Regulation of RNA biosynthetic process | 5,113 | 11 | 2.99 | 6.40E-05 |
| GO:0051252 | Regulation of RNA metabolic process | 5,162 | 11 | 3.02 | 7.00E-05 |
| GO:0019219 | Regulation of nucleobase-containing compound metabolic process | 5,317 | 11 | 3.11 | 9.10E-05 |
| GO:0048573 | Photoperiodism, flowering | 157 | 3 | 0.09 | 0.00011 |

**Table S5**: Selected genes regulated by both *PHYB* and *PHYC* in both SD and LD photoperiods. Fold change between WT and respective *phy*-null mutants are presented. The values for LD are the mean fold-change in expression between both experimental replicates. a - *FT-A1* and *FT-B1* expression was zero in the *phy*-null mutants in these comparisons.

| | | Fold change in expression | | | |
|---|---|---|---|---|---|
| | | Short days | | Long days | |
| **Gene ID** | **Annotation** | **WT/*phyB*-null** | **WT/*phyC*-null** | **WT/*phyB*-null** | **WT/*phyC*-null** |
| TraesCS7A01G115400 | *FT-A1* | - [a] | 168.88 | - [a] | - [a] |
| TraesCS7B01G013100 | *FT-B1* | - [a] | 286.72 | 927.23 | 481.46 |
| TraesCS3A01G143100 | *FT-A2* | 38.02 | 78.31 | 119.44 | 16.64 |
| TraesCS1B01G351100 | *FT-B3* | 14.44 | 14.18 | 3.56 | 7.52 |
| TraesCS5A01G391700 | *VRN-A1* | 3.09 | 2.66 | 15.64 | 7.96 |
| TraesCS5B01G396600 | *VRN-B1* | 70.61 | 52.50 | 113.15 | 73.89 |
| TraesCS2A01G261200 | *FUL-A2* | 92.75 | 644.50 | 800.56 | 127.00 |
| TraesCS2A01G174300 | *FUL-A3* | 351.75 | 369.78 | 740.74 | - |
| TraesCSU01G196100 | *PPD-B1* | 25.37 | 9.95 | 322.38 | 35.17 |
| TraesCS6A01G273200 | MYB | 21.12 | 8.30 | 43.09 | 2.89 |
| TraesCS6B01G300600 | MYB | 7.99 | 5.11 | 4.05 | 3.63 |
| TraesCS1A01G220300 | CONSTANS-like | 3.36 | 2.90 | 3.82 | 2.48 |
| TraesCS3B01G365300 | VQ-motif protein | 0.19 | 0.31 | 0.01 | 0.08 |
| TraesCS5A01G520200 | *TaFLC-A2* | 0.35 | 0.23 | 0.12 | 0.18 |
| TraesCS4B01G351500 | *TaFLC-B2* | 0.18 | 0.11 | 0.01 | 0.02 |
| TraesCS3A01G435000 | *TaFLC-A4-1* | 0.22 | 0.06 | 0.14 | 0.12 |
| TraesCS2A01G427200 | *WCOR15* | 0.03 | 0.11 | 0.01 | 0.05 |

**Table S6**: Selected genes regulated by both *PHYB* and *PHYC* in SD photoperiods but not LD photoperiods. Fold change between WT and respective *phy*-null mutants are presented as the mean fold-change between both experimental replicates. a - *FT-A4* expression was zero in the *phy*-null mutants in these comparisons.

| | | **Fold change in short days** | |
|---|---|---|---|
| **Gene ID** | **Annotation** | **WT/*phyB*-null** | **WT/*phyC*-null** |
| TraesCS3B01G162000 | *FT-B2* | 24.89 | 12.53 |
| TraesCS2A01G132300 | *FT-A4* | - [a] | 6.80 |
| TraesCS7B01G158900 | *TaFLC-B1* | 12.18 | 3.66 |
| TraesCS2B01G378700 | NF-YB | 2.70 | 3.07 |
| TraesCS7A01G233300 | MYB_related | 4.26 | 9.05 |
| TraesCS7B01G131600 | MYB_related | 14.02 | 16.90 |
| TraesCS5B01G054800 | bHLH | 5.55 | 2.59 |
| TraesCS5B01G183700 | WRKY | 7.80 | 5.16 |
| TraesCS3B01G129900 | WRKY | 13.83 | 6.20 |
| TraesCS3B01G240200 | WRKY | 17.62 | 6.85 |
| TraesCS7B01G249700 | WRKY | 123.51 | 205.05 |
| TraesCS3B01G199000 | WRKY | 2.38 | 2.94 |
| TraesCS5B01G183800 | WRKY | 4.22 | 2.92 |
| TraesCS6A01G146900 | WRKY | 4.96 | 2.58 |
| TraesCS1B01G374900 | WRKY | 5.07 | 2.47 |
| TraesCS3A01G347500 | WRKY | 5.51 | 2.87 |
| TraesCS1A01G334400 | GA-2oxidase | 0.05 | 0.13 |
| TraesCS6A01G143900 | BBox | 0.06 | 0.17 |
| TraesCS2A01G100600 | G2-like | 0.22 | 0.40 |
| TraesCS3A01G274900 | GATA | 0.55 | 0.46 |
| TraesCS1B01G335100 | CBF | 0.16 | 0.17 |
| TraesCS5A01G311100 | CBF | 0.19 | 0.37 |
| TraesCS5A01G310800 | CBF | 0.47 | 0.18 |
| TraesCS5A01G310700 | CBF | 0.52 | 0.33 |

**Table S7**: Selected genes regulated by both *PHYB* and *PHYC* in LD photoperiods but not SD photoperiods. Fold change between WT and respective *phy*-null mutants are presented as the mean fold-change between both experimental replicates.

| | | **Fold change in long days** | |
|---|---|---|---|
| **Gene ID** | **Annotation** | **WT/*phyB*-null** | **WT/*phyC*-null** |
| TraesCS3A01G116300 | *GIGANTEA-A* | 2.35 | 3.21 |
| TraesCS3B01G135400 | *GIGANTEA-B* | 2.59 | 3.99 |
| TraesCS6B01G315400 | CO-like | 0.42 | 0.23 |
| TraesCS7B01G115200 | SPL | 0.24 | 0.25 |
| TraesCS3A01G107200 | *TZF-A1* | 0.09 | 0.18 |

**Table S8:** *PHYB*- and *PHYC*-regulated splicing-related genes. Mean TPM and fold-change for each genotype was calculated from four biological replicates in SD samples, and eight biological replicates for LD samples.

| ***PHYB* regulated under SD** | | **TPM** | | |
|---|---|---|---|---|
| **Gene ID** | **Annotation** | **Wild-type SD** | ***phyB*-null_SD** | **Fold-change** |
| TraesCS3B01G157400 | Pre-mRNA-splicing factor of RES complex protein | 6.4 | 3.4 | 1.9 |
| TraesCS1B01G125800 | Pre-mRNA-splicing factor cwc26 | 10.0 | 2.3 | 4.3 |
| TraesCS7B01G194600 | Pre-mRNA-splicing factor SLU7 | 6.6 | 3.3 | 2.0 |
| TraesCS5B01G017300 | Splicing factor 3B subunit 1 | 0.2 | 0.9 | 0.2 |
| TraesCS7B01G450300LC | Splicing factor 3B subunit 4 | 0.0 | 2.0 | - |
| TraesCS2B01G720700LC | Splicing factor U2AF small subunit A | 0.1 | 3.2 | 0.0 |
| TraesCS4B01G209600 | SR-rich pre-mRNA splicing activator | 4.4 | 1.7 | 2.6 |
| TraesCS4A01G094900 | SR-rich pre-mRNA splicing activator | 25.2 | 4.9 | 5.1 |
| TraesCS2A01G122400 | Splicing factor U2AF, large subunit | 3.0 | 1.3 | 2.3 |
| TraesCS7A01G075100 | Splicing factor U2AF large subunit A | 0.2 | 0.0 | 17.9 |
| ***PHYC*-regulated under SD** | | **TPM** | | |
| **Gene ID** | **Annotation** | **Wild-type SD** | ***phyC*-null SD** | **Fold-change** |
| TraesCS1B01G125800 | Pre-mRNA-splicing factor cwc26 | 10.0 | 2.2 | 4.6 |
| TraesCS2A01G646800LC | Pre-mRNA-processing-splicing factor 8 | 0.1 | 1.1 | 0.1 |
| TraesCS1B01G416400 | Pre-mRNA-processing-splicing factor 8 | 3.1 | 0.2 | 15.6 |
| TraesCS1B01G130200 | Arginine/serine-rich splicing factor | 0.2 | 1.8 | 0.1 |
| TraesCS5B01G017300 | Splicing factor 3B subunit 1 | 0.2 | 0.6 | 0.3 |
| TraesCS7B01G450300LC | Splicing factor 3B subunit 4 | 0.0 | 2.7 | - |
| TraesCS7B01G384700LC | Splicing factor family-like | 0.5 | 0.0 | - |
| TraesCS5A01G366000 | Splicing factor U2AF large subunit | 0.0 | 0.2 | 0.2 |
| TraesCS2B01G720700LC | Splicing factor U2AF small subunit A | 0.1 | 2.1 | 0.0 |
| ***PHYB*-regulated under LD** | | **TPM** | | |
| **Gene ID** | **Annotation** | **Wild-type LD** | ***phyB*-null LD** | **Fold-change** |
| TraesCS2A01G122400 | Splicing factor U2AF, large subunit | 3.5 | 1.5 | 2.3 |
| TraesCS3A01G086900 | Splicing factor | 5.8 | 9.8 | 0.6 |
| TraesCS4A01G399500 | Pre-mRNA-splicing factor CWC22-like protein | 3.0 | 9.1 | 0.3 |
| TraesCS3A01G426500 | Lysine ketoglutarate reductase trans-splicing-like protein | 14.2 | 7.0 | 2.0 |
| TraesCS5A01G189600 | Pre-mRNA-processing-splicing factor 8 | 0.9 | 1.6 | 0.6 |
| TraesCS3B01G293100 | Arginine/serine-rich splicing factor | 9.3 | 11.6 | 0.8 |
| TraesCS3A01G260000 | Arginine/serine-rich splicing factor | 11.7 | 18.4 | 0.6 |

| | | | | |
|---|---|---|---|---|
| TraesCS4A01G091300 | RNA-binding protein | 37.6 | 53.9 | 0.7 |
| TraesCS1B01G244200 | Splicing factor-like protein | 4.3 | 6.5 | 0.7 |
| TraesCS5A01G267500 | Splicing factor U2AF small subunit A | 19.9 | 34.7 | 0.6 |
| ***PHYC*-regulated under LD** | | **TPM** | | |
| **Gene ID** | **Annotation** | **Wild-type LD** | ***phyC*-null LD** | **Fold-change** |
| TraesCS1B01G130200 | Arginine/serine-rich splicing factor | 0.1 | 0.5 | 0.2 |
| TraesCS1B01G615200LC | Splicing factor 3B subunit 5 | 2.1 | 11.7 | 0.2 |

**Table S9:** Top 40 significant enriched GO terms among genes impacted by any AS event by either *PHYB* or *PHYC* under SD or LD conditions.

| GO ID | Term | Annotated | Significant | Expected | *P* |
|---|---|---|---|---|---|
| GO:0009662 | Etioplast organization | 5 | 3 | 0.05 | 1.20E-05 |
| GO:0006400 | tRNA modification | 204 | 11 | 2.17 | 1.40E-05 |
| GO:0019988 | Charged-tRNA amino acid modification | 7 | 3 | 0.07 | 4.10E-05 |
| GO:0008033 | tRNA processing | 328 | 12 | 3.49 | 0.00024 |
| GO:0030488 | tRNA methylation | 61 | 5 | 0.65 | 0.00049 |
| GO:0044255 | Cellular lipid metabolic process | 1,877 | 36 | 20 | 0.00056 |
| GO:0030258 | Lipid modification | 287 | 10 | 3.06 | 0.00112 |
| GO:0015855 | Pyrimidine nucleobase transport | 6 | 2 | 0.06 | 0.00165 |
| GO:0015857 | Uracil transport | 6 | 2 | 0.06 | 0.00165 |
| GO:0035344 | Hypoxanthine transport | 6 | 2 | 0.06 | 0.00165 |
| GO:0098702 | Adenine import across plasma membrane | 6 | 2 | 0.06 | 0.00165 |
| GO:0098710 | Guanine import across plasma membrane | 6 | 2 | 0.06 | 0.00165 |
| GO:0098721 | Uracil import across plasma membrane | 6 | 2 | 0.06 | 0.00165 |
| GO:1903716 | Guanine transmembrane transport | 6 | 2 | 0.06 | 0.00165 |
| GO:1903791 | Uracil transmembrane transport | 6 | 2 | 0.06 | 0.00165 |
| GO:1904082 | Pyrimidine nucleobase transmembrane transport | 6 | 2 | 0.06 | 0.00165 |
| GO:0046856 | Phosphatidylinositol dephosphorylation | 48 | 4 | 0.51 | 0.00171 |
| GO:0043412 | Macromolecule modification | 9,166 | 124 | 97.66 | 0.00191 |
| GO:0006650 | Glycerophospholipid metabolic process | 476 | 13 | 5.07 | 0.002 |
| GO:0006629 | Lipid metabolic process | 2,471 | 42 | 26.33 | 0.00207 |
| GO:0010167 | Response to nitrate | 27 | 3 | 0.29 | 0.00291 |
| GO:0015853 | Adenine transport | 8 | 2 | 0.09 | 0.00304 |
| GO:0015854 | Guanine transport | 8 | 2 | 0.09 | 0.00304 |
| GO:0072531 | Pyrimidine-containing compound transmembrane transport | 8 | 2 | 0.09 | 0.00304 |
| GO:0006560 | Proline metabolic process | 29 | 3 | 0.31 | 0.00357 |
| GO:0046488 | Phosphatidylinositol metabolic process | 340 | 10 | 3.62 | 0.00382 |
| GO:0006464 | Cellular protein modification process | 8,493 | 114 | 90.49 | 0.00401 |
| GO:0036211 | Protein modification process | 8,493 | 114 | 90.49 | 0.00401 |
| GO:0006399 | tRNA metabolic process | 709 | 16 | 7.55 | 0.00434 |
| GO:0046486 | Glycerolipid metabolic process | 530 | 13 | 5.65 | 0.00495 |
| GO:0046839 | Phospholipid dephosphorylation | 65 | 4 | 0.69 | 0.00516 |
| GO:0098739 | Import across plasma membrane | 11 | 2 | 0.12 | 0.00585 |
| GO:0006396 | RNA processing | 1,795 | 31 | 19.13 | 0.00619 |
| GO:0051179 | Localization | 6,909 | 94 | 73.62 | 0.00663 |
| GO:0001510 | RNA methylation | 259 | 8 | 2.76 | 0.00696 |
| GO:0006282 | Regulation of DNA repair | 12 | 2 | 0.13 | 0.00697 |

| | | | | | |
|---|---|---|---|---|---|
| GO:1904823 | Purine nucleobase transmembrane transport | 12 | 2 | 0.13 | 0.00697 |
| GO:0006810 | Transport | 6,587 | 90 | 70.18 | 0.00712 |
| GO:0009451 | RNA modification | 624 | 14 | 6.65 | 0.00772 |
| GO:2001020 | Regulation of response to DNA damage stimulus | 13 | 2 | 0.14 | 0.00818 |
